# Direct and indirect fitness effects of plant metabolites and genetic constraints limit evolution of allelopathy in an invading plant

**DOI:** 10.1101/2023.06.27.546761

**Authors:** Richard Honor, Mia Marcellus, Robert I. Colautti

**Affiliations:** Queen’s University, 99 University Ave, Kingston, ON K7L 3N6

## Abstract

1. Invading species encounter novel communities of consumers, pathogens, and competitors. Both phenotypic plasticity and rapid evolution can facilitate invasion across these heterogenous communities. However, the rate and extent of adaptive evolution on contemporary timescales can be constrained by adaptive phenotypic plasticity and the genetic architecture of traits under selection.
2. We measured phenotypic plasticity and quantified genetic variation for growth, leaf chlorophyll *a* (Chl *a*) and glucosinolates, and lifetime fitness among 23 naturally inbred seed families of *Alliaria petiolata* (garlic mustard) collected across its invasive range in eastern North America. After growing a self-pollinated generation in a uniform common garden to reduce maternal effects, we reared second-generation plants in a two-year greenhouse and field experiment with naïve soil from an uninvaded habitat. We estimated selection gradients and causal factors affecting lifetime fitness when reared alone, with an intraspecific competitor, and under interspecific competition with naïve *Acer saccharum* (sugar maple) saplings.
3. We defined Total Metabolite Production (TMP) as the first principal component of Chl *a* and glucosinolate concentrations, accounting for 84% of variation in these two traits. TMP was significantly plastic across growing environments (*p* < 0.001) with limited broad-sense heritability (*H^2^* = 2.91; *p* = 0.08). Path analysis revealed that plastic phenotypes with higher TMP had an indirect positive effect on *A. petiolata* fitness via a direct, negative effect on performance of *A. saccharum* competitors. In contrast, the second principal component defined Relative Glucosinolate Investment (RGI), which was significantly heritable (*H^2^* = 16.91, p < 0.001) with no detectable plasticity across treatments. Variation in RGI among *A. petiolata* genotypes had a direct, positive effect on *A. saccharum* performance and an indirect negative effect on *A. petiolata* fitness.
4. *Synthesis.* Adaptive evolution of allelopathy during invasion has been constrained by (i) a lack of heritable genetic variation for allelopathy, (ii) high plasticity for TMP across competition treatments, and (iii) selection for lower RGI under interspecific and intraspecific competition. As an alternative to eco-evolutionary feedbacks, plasticity in TMP may be an overlooked explanation for variable performance of *A. petiolata* across its introduced range.

## Introduction

Invasive species, from a broad range of taxa, have invaded new geographic regions and out-performed native species despite having had less time to adapt to local conditions. To explain this apparent paradox, several hypotheses have proposed rapid, adaptive evolutionary responses to novel biotic interactions. For example, the Evolution of Increased Competitive Ability (EICA) hypothesis posits that the loss of specialist herbivores favours the evolution of more competitive genotypes when costly defences are no longer needed (Blossey and Notzold 1995; Keane and Crawley 2002; Müller-Schärer et al. 2004; Doorduin and Vrieling 2011). The Novel Weapons Hypothesis (NWH) attributes the dominance of invasive species to allelopathic metabolites that suppress native competitors (Inderjit et al. 2008; Gomes et al. 2017; Hierro and Callaway 2021; Inderjit et al. 2021). The roles of natural enemies and competition are not mutually exclusive; for example, a reduction in herbivory could favour an evolutionary shift in chemistry from defensive to allelopathic (Uesugi and Kessler 2013). Genetic changes predicted by these hypotheses are typically framed within a dichotomy between native and introduced ranges but could equally apply across introduced populations to explain variability in invasion rates. Indeed, there is evidence that ecologically relevant variation among introduced populations is often overlooked, acting as lurking variables in comparative studies framed within the native-introduced dichotomy (Colautti et al. 2009; Colautti and Lau 2015). Evolutionary responses to biotic interactions that differ across the introduced range could facilitate the establishment and spread of invasive species, and account for variability in invasion rates throughout the introduced range.

There is some evidence that eco-evolutionary feedbacks within the introduced range can facilitate invasion. Greenhouse and growth chamber experiments demonstrate how invasions can favour life history traits at the expanding invasion front, which in turn feed back to increase rates of spread (Williams et al. 2016; Ochocki and Miller 2017; Szücs et al. 2017). In Australia, *Rhinella marina* (cane toads) evolved morphological changes that increase dispersal at the invasion front, facilitating further spread (Philips et al. 2006; Shine et al. 2021). In North America, the invasive perennial herb *Lythrum salicaria* (purple loosestrife) rapidly evolved clines in flowering time (Wu and Colautti 2022) as an adaptive response to local season length, which in turn increased reproductive output by an order of magnitude (Colautti and Barrett 2013). These examples demonstrate how evolution can occur rapidly along abiotic environments and density gradients, resulting in considerable variation in genotype, phenotypes, and population growth rates across the introduced range. However, less is known about the eco-evolutionary response of invasive species to biotic interactions.

Methods for the study of evolution in natural populations have been refined for decades (Endler 1986; Lande and Arnold 1983), yet these are rarely applied to studies of biological invasions (Colautti and Lau 2015). The ‘Chicago School’ of ecological genetics separates natural selection, which acts on phenotypes, from heritable genetic variation, which is necessary for an evolutionary response to selection (Endler 1986; Conner and Hartl 2004; Svennson 2023). Moreover, the evolution of highly heritable traits under strong selection can be constrained nonetheless by trait correlations. For example, chlorophyll *a* (Chl *a*) and glucosinolate production are often correlated due to pleiotropy (Chen et al. 2012) and chromosomal linkage (Qian et al. 2016). Additionally, the magnitude and direction of natural selection are environment-dependent, potentially mediated by biotic interactions that are not easily assessed by selection gradients alone (Scheiner et al. 2000). Instead, path analysis, structural equation models, and other forms of causal analysis can test hypotheses about the biotic context of natural selection acting through direct and indirect mechanisms (Kingsolver and Schemske 1991; Scheiner et al. 2000; Shipley 2016). For example, the NWH implicitly predicts that allelochemicals increase fitness indirectly via direct suppression of competitors. When used in concert, selection gradients, quantitative genetics, and causal analysis provide a versatile toolkit for investigating eco-evolutionary dynamics of biological invasions.

Glucosinolate concentration in *Alliaria petiolata* (garlic mustard) is a good model trait for investigating environment-dependent, eco-evolutionary dynamics. Like many invasive species, populations of *A*. *petiolata* shift from primarily interspecific competition toward intraspecific competition as populations grow. In *A. petiolata*, it has been proposed that an eco-evolutionary feedback between glucosinolate production and competition select for genetic populations that differ in glucosinolate production, depending on the ratio of intra- to inter-specific competition (Lankau et al. 2009; Evans et al. 2016). Although glucosinolate production is environmentally induced and maternally inherited (Holeski et al. 2012), previous studies have not accounted for maternal effects, nor measure plasticity, heritability, or natural selection for glucosinolates. Many experiments have tested the allelopathic effects of glucosinolates, but mostly in controlled environments and with mixed conclusions about their role in *A*. *petiolata* invasion (Roberts and Anderson 2001; Stinson et al. 2006; Burke 2008; Callaway et al. 2008; Wolfe et al. 2008; Barto and Cipollini 2009; Barto et al. 2011; Cantor et al. 2011; Burke et al. 2019; Duchesneau et al. 2021; Roche et al. 2021). Therefore, our goal was to experimentally remove maternal effects and test predictions of the eco-evolutionary feedbacks model using selection gradients, quantitative genetics, and causal analysis.

Here, we test the hypothesis that introduced populations of *A. petiolata* evolved local differences in glucosinolate investment as expanding populations shifted from inter- to intra-specific competition. First, we use mixed models to assess genetic variation and plasticity for glucosinolate investment, growth, and fitness in *A*. *petiolata* genotypes sampled across eastern North America. Genetic divergence among populations is the main predicted outcome of eco-evolutionary feedbacks. An alternative hypothesis is that adaptive phenotypic plasticity limits adaptive evolution yet facilitates invasion across heterogenous environments. Second, we analyze selection gradients to test whether natural selection favours higher glucosinolate investment under interspecific competition and lower investment under intraspecific competition or absent competition. This fluctuating selection is required for a negative feedback to drive eco-evolutionary dynamics. Finally, we test mechanistic predictions of ecological feedbacks using path analysis. Specifically, we quantify the direct effects of glucosinolate production by *A*. *petiolata* on growth of competing *Acer saccharum* (sugar maple), and the resulting indirect effect on *A*. *petiolata* fitness. This tests the key prediction that natural selection favours higher glucosinolate production indirectly by suppressing competition.

## Materials and Methods

### Seed sources

*Alliaria petiolata* seeds were collected between 2009–2012 as part of the Global Garlic Mustard Field Survey (Colautti et al. 2014) with methods detailed therein. Of the 229 sample locations across North America, 23 populations with viable seeds were chosen to maximize the geographic scope and range of ecological conditions among introduced populations, including invasion stage, density, herbivory, pathogen damage, and habitat. Our experiment uses the offspring of a single plant per population (i.e., “seed family”) as a representative of each naturally inbred parent population. Seed families are thought to adequately capture the genetic profile of the parent population because *A. petiolata* populations are highly inbred, with molecular markers showing little genetic variation within populations (Durka et al. 2005). To dilute maternal effects on growth and fitness, one individual from each family was first grown in an outdoor common garden at the Queen’s University Biological Station (QUBS; 44.5671° N, 76.3250° W) in 2017-2018, and seeds produced from these offspring were used in the experiment. Four of the 23 seed families did not produce viable seeds, and were replaced with seeds grown from the original field collection (Figure 1; Table 1). This includes one genotype from the native range (EFCC2-4-80), which was included because it is from a source population close to the genotype used in the garlic mustard genome project (Alabi et al. 2021). However, excluding it from the final analysis did not appreciably change any of the results presented.

**Figure 1.**
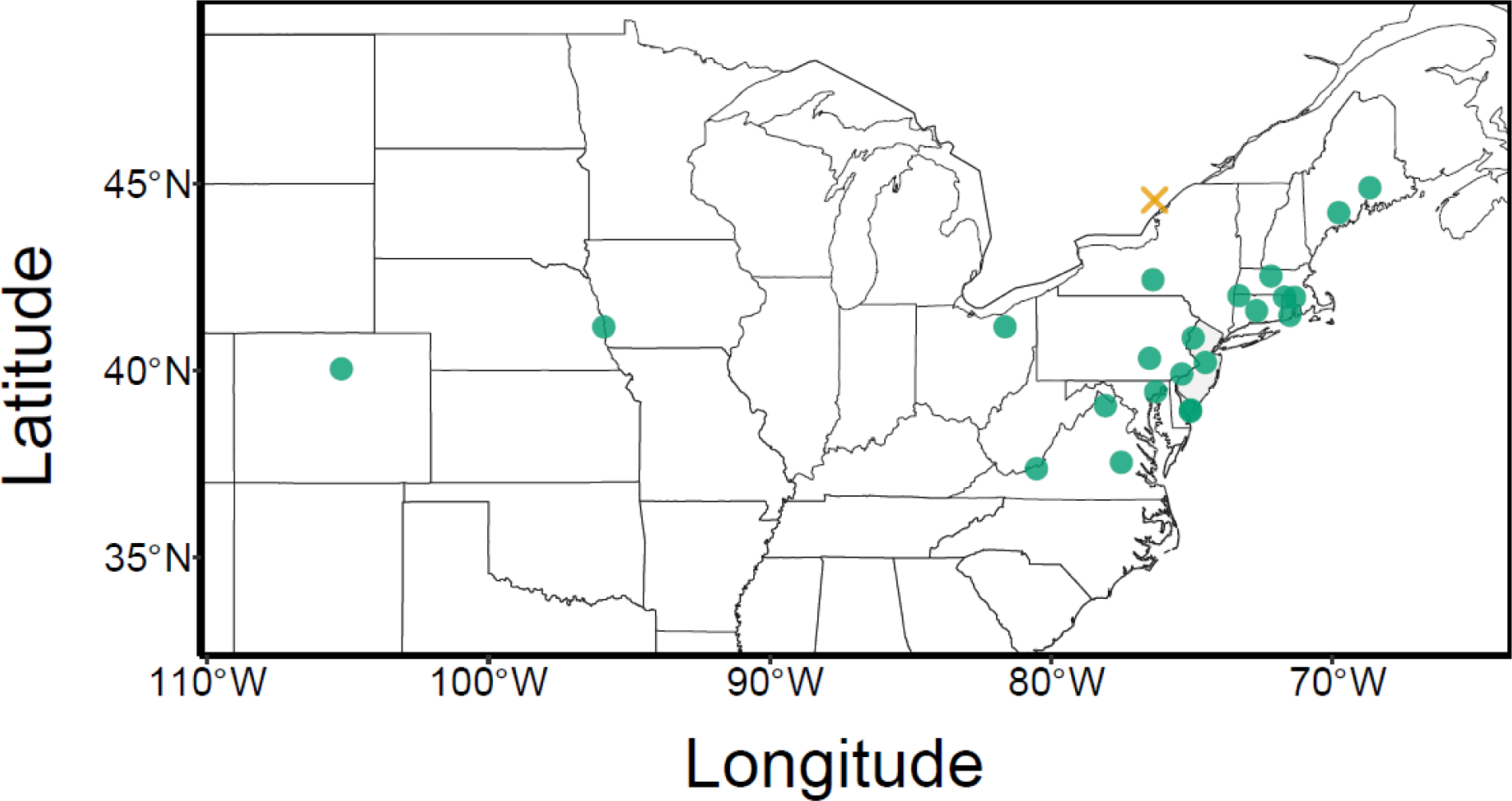
Sampling locations of naturally inbred seed families used in the experiment (green dots) excluding one family (EFCC2-4-80), which was sampled from northern Italy. The yellow “X” indicates the location of the field common garden site at the Queen’s University Biological Station (QUBS), and where *Acer saccharum* seedlings were collected.

**Table 1.** Summary of seed sampling locations and number of surviving individuals from each accession in each of three treatments: alone, intraspecific competition, and interspecific competition.

| Accession | Lat | Long | Alt (m) | N Alone | N Intra | N Inter |
| --- | --- | --- | --- | --- | --- | --- |
| JBCHY1-1-50 | 38.95 | -75.05 | 25.9 | 8 | 9 | 9 |
| JBBLB2-1-20 | 38.91 | -75.05 | 0 | 7 | 7 | 8 |
| CRSOSO-3-40 | 44.9 | -68.68 | 59 | 8 | 6 | 7 |
| SMITH1-20 | 42.43 | -76.39 | 326 | 4 | 9 | 8 |
| CWRIC2-X2 | 37.54 | -77.51 | 30 | 8 | 5 | 7 |
| DVGM-20 | 40.88 | -74.96 | 232 | 7 | 5 | 8 |
| KVEDG1-1-80 | 39.44 | -76.29 | 2 | 6 | 8 | 6 |
| MHBUR1-4-20 | 41.97 | -71.71 | 128 | 8 | 5 | 7 |
| WSSWM3-1-0 | 39.91 | -75.36 | 23 | 6 | 5 | 9 |
| JWBOY-2-80 | 39.06 | -78.07 | 250.9 | 7 | 6 | 5 |
| MAVBEL2-1-20* | 41.17 | -95.9 | 358 | 5 | 6 | 6 |
| VRCAN-1-40 | 42 | -73.33 | 203 | 6 | 4 | 6 |
| VRPET2-3-20 | 42.53 | -72.19 | 329.1 | 8 | 4 | 4 |
| BWPEM1-9-0 | 37.37 | -80.52 | 1156 | 4 | 6 | 5 |
| EFCC2-4-80 | 45.7 | 11.69 | 73 | 4 | 4 | 5 |
| JARI1-1-80 | 41.48 | -71.52 | 20.7 | 5 | 5 | 3 |
| SMAKC1-B* | 41.17 | -81.64 | 307 | 4 | 4 | 5 |
| VSGARO1-1-0 | 44.23 | -69.78 | 62 | 5 | 3 | 5 |
| MSMID1-1-20 | 41.6 | -72.7 | 5 | 4 | 2 | 4 |
| CBMCK1-1-0* | 40.04 | -105.23 | 1606.9 | 4 | 4 | 1 |
| PDVRT1-20 | 40.22 | -74.52 | 52 | 3 | 4 | 2 |
| MHNAT1-2-0 | 41.96 | -71.35 | 60 | 2 | 2 | 3 |
| RULEB1-20* | 40.33 | -76.51 | 146.3 | 2 | 3 | 1 |
| **Total** |  |  |  | **125** | **116** | **124** |

\*Seed families were derived from the original field collection

On May 8, 2019, we collected *Acer saccharum* saplings from a natural population located in a forest understory adjacent to a gravel road near the common garden site at QUBS. This section of the forest was not yet invaded, but it appeared to be suitable habitat as there was a recently established (< 5 y) and expanding population of *A. petiolata* nearby (∼ 25 m). Blocks of soil, ∼15 cm deep with an abundance of *A. saccharum* saplings, were transported back to the greenhouse where saplings were placed into experimental pots. We retained soil around roots to avoid disrupting root hairs and associated mycorrhizal connections. Additional field soil was collected from this area by removing debris and other plant material from the surface of the soil, and then collecting the top layer to a depth of ∼ 12 cm. The soil was then thoroughly homogenized using shovels and rakes prior to use in the glasshouse.

### Experimental design

To limit human error in labelling and data collection, we used the baRcodeR package in R to generate a unique, scannable 2-dimensional barcode for each plant (Wu et al. 2020). As detailed in Appendix A of the Supporting Information, we stratified seeds of *Alliaria petiolata* in petri dishes and transferred germinated seedlings into 2cm x 2cm plug trays at a constant 10°C (12h half daylight) in a growth chamber to limit growth of early-germinating seeds. On May 9-10, 2019, we transplanted all available seedlings into 4-inch plastic pots in a glasshouse at the Queen’s University Phytotron under natural light, filtered with a shade cloth and watered as needed. Sample size varied across 23 families due to variation in germination, survival, and sample loss associated with glucosinolate and Chl *a* processing. As shown in Table 1, the effective total sample size was 365 *A. petiolata* plants, with a median of 16 plants per seed family.

Starting with 23 genetically distinct *A. petiolata* accessions from the Global Garlic Mustard Field Survey (Colautti et al. 2014), we grew at least 3 individuals from each accession in each of 3 different competitive environments: (i) alone, (ii) interspecific competition with *A. saccharum*, and (iii) intraspecific competition with a different *A. petiolata* seed family (Table 1; Supporting Information Figure A1). We also grew 31 *A. saccharum* seedlings alone (i.e., the “maple alone” treatment), as a control for comparison with *A. saccharum* growth and survival in the interspecific treatment. In natural conditions, *A. petiolata* density can be as many as 18 plants/dm^2^ in April and dwindle to as little as 0.6 plants/dm^2^ by June (Anderson et al. 1996). In comparison, density in our experiment was either 1.23 plants/dm^2^ in the alone treatment or 2.46 plants/dm^2^ in the intraspecific treatment.

In the first growing season, plants were kept in the Queen’s University Phytotron to limit environmental influences on leaf chemistry traits, growth rate, and total leaf area. *Alliaria petiolata* is a biennial herb that requires an overwinter vernalization period to promote flowering (Cavers et al. 1979). Therefore, plants were transplanted from the glasshouse into a field site (QUBS, 44.56 °N, 76.32 °W) at the end of the first growing season, to allow plants to acclimatize to cold temperatures before winter. This not only promoted flowering of *A. petiolata*, but additionally allowed us to assess overwinter survival and lifetime fecundity under natural field conditions.

### Field common garden

To allow plants to acclimate to colder temperatures, on October 5, 2019, we transplanted all surviving plants from the greenhouse into a field plot at the QUBS field site, which included 2 m tall aluminum fencing to exclude vertebrate herbivores. To limit interactions with surrounding vegetation, we buried pots directly into the field soil, arranged into 49 experimental blocks with dimensions 0.6 m × 1.2 m and 15 cm deep, with pots spaced 10 cm apart.

### Greenhouse measurements (year 1)

To account for variation in time to germination and seedling size at transplant, we measured the length of the largest leaf and the number of true leaves on *A. petiolata* on May 16 and 17, 2019. To account for variation in *A. saccharum* size at transplant, we measured stem height, the number of true leaves and the length of the largest leaf on *A. saccharum*. At the end of the first growing period in the glasshouse (August 14 to 21, 2019), we estimated total leaf area (TLA) on both plants as the leaf length multiplied by leaf width, summed across all leaves (hereafter TLA_AS_ for *A. saccharum* and TLA_AP_ for *A. petiolata*).

Growth rates of *A. petiolata* were calculated from rosette growth in bi-weekly measurements. As detailed in Appendix A of the Supporting Information, we used computer imaging with a machine learning algorithm, validated by hand-measurements, to determine the total leaf area of each plant in the experiment.

We quantified Chl *a* and total glucosinolate concentration in *A. petiolata* using methods described in Appendix A of the Supporting Information. Sinigrin was used for the glucosinolate standard curve as it is one of the most abundant glucosinolates in *A. petiolata* and it is suspected to have allelopathic effects (Vaughn and Berhow 1999).

### Field measurements (year 2)

On May 13, 2020, we recorded overwinter survivorship. On July 8 and 9, 2020, we collected the siliques from all remaining *A. petiolata* plants, and the above-ground shoots of all remaining *A. saccharum* plants. All plant material was then dried to constant weight. We measured silique biomass as our estimate of reproductive fitness in *A. petiolata*, and shoot biomass as our estimate of *A. saccharum* performance. As a biennial, lifetime fitness of *A. petiolata* was calculated as the joint distribution of survival and silique biomass at the end of the experiment, including plants that did not survive to reproduce (lifetime fitness = 0).

### Biotic interactions

By using non-sterile field soil, we allowed plants to be exposed to pathogens and other organisms in the soil. We also allowed germination of *Sphagnum* moss, which covered the soil surface, and ferns from the family Dryopteridaceae, which colonized sporadically. On October 1, 2019, the number of ferns in each pot were counted to a maximum of 15 individual plants to use as a covariate in our analysis. On *A. petiolata* leaves, we observed thrips (order Thysanoptera), a powdery mildew fungus identified as *Erysiphe cruciferarum*, and black rot (*Xanthomonas campestris*). These pathogens commonly infect the Brassicaceae (Ciola and Cipollini 2011; Vicente and Holub 2013) and were identified by characteristics leaf lesions (see Supporting Information Figure A2 for examples).

Enemy abundance was evaluated by photographing the same leaves that were chosen for glucosinolate and Chl *a* quantification. On each leaf, thrip damage and *X. campestris* lesions were counted, and the presence of *E. cruciferarum* was recorded. *Acer saccharum* also exhibited leaf damage, observable as leaf holes likely of thrip origin, which were small but still visible to the naked eye. The presence of leaf damage on each leaf was recorded while TLA_AS_ was measured, and the proportion of damaged *A. saccharum* leaves on each plant was used to indicate pathogen damage for that species.

### Data analysis

To account for an expected correlation between Chl *a* and total glucosinolate concentration, we used a principal component analysis (PCA) to identify two orthogonal yet biologically meaningful principal component ‘traits.’ We refer to the first principle component as Total Metabolite Production (TMP) because it correlates with both Chl *a* and total glucosinolate concentration, as shown later in the Results; we refer to the second principle compent as Relative Glucosinolate Investment (RGI), because it measures the concentration of glucosinolates relative to Chl *a*. A similar approach was used for phenological traits (Colautti and Barrett 2010) because PCA can address statistical complications involving collinearity among biological traits (Chong et al. 2018). In addition to the PCA on Chl *a* and total glucosinolate concentration, we used two additional PCA to summarize the transplant size measurements of *A. saccharum* (TS_AS_) and *A. petiolata* (TS_AP_), where TS_AS_ and TS_AP_ are the first principal component axis of each analysis (see Supporting Information).

We used statistical mixed models to estimate the proportion of total phenotypic variation attributable to genetic variation (G) among inbred seed families of *A*. *petiolata*, plasticity due to environmental variation across competition treatments (E), and genetic variation for plasticity (G × E). Separate models for each *A. petiolata* phenotypic trait as a dependent variable included seed family, treatment, and a family-by-treatment interaction as random effects. Mixed effects models were fit to the data using the package glmmTMB in R (Brooks et al. 2017; R Core Team 2020) and the significance of each effect was assessed using a likelihood ratio test. Greenhouse bench and position within the field common garden were included only when significant, and we included leaf area as a covariate in models of leaf chemistry traits. Reproducible R code for all statistical models is available, along with all original data, in a repository archived through Dryad (https://doi.org/10.5061/dryad.cfxpnvxbj).

### Selection analysis

We measured selection using standardized selection gradients (Lande and Arnold 1983; Matsumura et al. 2012). Mixed effects models were fitted to four separate response variables: TLA_AP_, survival, fecundity, and lifetime fitness. Each full model included an interaction term for TMP and treatment, and a second interaction term between RGI and treatment. These two-way interactions represent a statistical test of whether selection on TMP and RGI differ across competition treatments. Our statistical models also included the fixed effects of bench, fern abundance, TS_AP_, relative growth rate, thrip damage, *X. campestris* infection, and *E. cruciferarum* presence. The random effect of seed family was also included in all maximum models. Fitness measured in the field common garden also included TLA_AP_ as a fixed effect (covariate) and the random effects of row and column describing the spatial location of each individual. A model of best fit for each fitness measurement was determined by sequentially performing likelihood ratio tests of hierarchical models derived from the full model outlined above.

A generalized linear model with a binomial error distribution and a logit link was used for model selection of survival rates. Minimum adequate models were then converted to standardized selection gradients after all terms were assessed for significance. Standardized selection gradients of survival data were obtained by dividing data by the mean survival rate and fitting the model with a gaussian error distribution (Hajduk et al. 2020). We did this because the predicted response to selection is proportional to the additive genetic covariance between a trait and fitness.

Selection gradients measure direct selection on a trait while controlling for selection on correlated traits. To calculate the change in mean trait value from one generation to the next, we measured selection differentials for TMP and RGI in each treatment by subtracting the phenotypic mean of all individuals from the phenotypic mean of individuals that survived; this estimated change in mean phenotype, which is a consequence of both direct selection on the trait and indirect selection on correlated traits.

As with *A. petiolata*, we used model selection with likelihood ratio tests of nested models to assess the significance of interspecific competition on TLA_AS_ in year 1 and on the log-transformed *A. saccharum* shoot biomass in year two (to meet normality assumptions). To assess the average effect of interspecific competition on *A. saccharum*, the fixed effects of TS_AS_, fern abundance, bench, leaf damage and treatment (i.e., *A. saccharum* alone vs interspecific competition treatment) were included in our statistical models.

We did not directly measure soil microbial communities or root exudates. To test whether these or other, unmeasured *A. petiolata* traits might affect *A. saccharum* performance, we again used model selection with likelihood ratio tests on *A. saccharum* performance in year 1 and 2. Fixed effects included TS_AS_, leaf damage and fern abundance, as well as the *A. petiolata* traits: relative growth rate, TLA_AP_, TMP, RGI, and seed family. Random effects included greenhouse bench, as well as field row and column in the year 2 model.

### Path analysis

As with selection gradients, path coefficients are analogous to partial regression coefficients but the latter can measure selection on a trait through hypothesized causal relationships (Scheiner et al. 2000). We used standardized path gradients to measure direct selection on glucosinolate production (TMP and RGI) resulting from the allelopathic effect of glucosinolates on *A. saccharum.* Four hypothetical path scenarios were analyzed, one for each *A. petiolata* performance/fitness measure as the response, namely TLA_AP_, survival, fecundity, and lifetime fitness. Each path scenario included the direct effects of TMP and RGI on *A. petiolata* performance/fitness, and the indirect effects of TMP and RGI on *A. petiolata* performance/fitness via impacts on *A. saccharum* performance in year 1. We also included the significant predictors of *A. saccharum* and *A. petiolata* performance in each. Greenhouse bench, field row and column were not included in path analyses, as path analysis cannot handle categorical variables. We used the Lavaan package (Rosseel 2012) for each path analysis, using permutation tests with 10,000 iterations to test for model significance.

## Results

Mortality in the greenhouse (year 1) was 2.29 % for *A. petiolata* and 1.53 % for *A. saccharum* but increased under field conditions (year 2) to 69.66 % for *A. petiolata* and 6.25 % for *A. saccharum*. Total leaf area of *A. petiolata* (TLA_AP_), fecundity, and lifetime fitness were lowest in the intraspecific treatment (Supporting Information Table B3). For example, mean fecundity was 3.13 g (95 % CI = ± 0.84 g) in the alone treatment, 1.78 g (95 % CI = ± 0.83 g) in the interspecific treatment and 1.07 g (95 % CI = ± 0.38 g) in the intraspecific treatment. In contrast, *A. petiolata* mortality was highest in the interspecific treatment (73.62 %) compared to 68.18 % (120/176) in the intraspecific treatment, and 65.27 % (109/167) in the alone treatment (Table B3).

### Metabolite correlations

Consistent with studies of other species, we found that concentrations of glucosinolate and Chl *a* were strongly correlated (r^2^ = 0.43, *p* < 0.001). In the PCA of glucosinolate and Chl *a* production, Total Metabolite Production (TMP) accounted for 84% of the variation in these two traits, whereas Relative Glucosinolate Investment (RGI) accounted for 16 % (Figure 2; Supporting Information Table B4). Higher TMP values represent individuals with greater leaf tissue concentrations of both glucosinolates and Chl *a*. In contrast, higher RGI values represent individuals with a higher production of glucosinolates relative to Chl *a*. Mean TMP was greatest in the alone treatment (0.47; 95 % CI = ±0.19), followed by the intraspecific treatment (–0.05; 95 % CI = ±0.26), and the interspecific treatment (–0.43; 95 % CI = ±0.21), while RGI was similar across treatments (Supporting Information Table B3).

**Figure 2.**
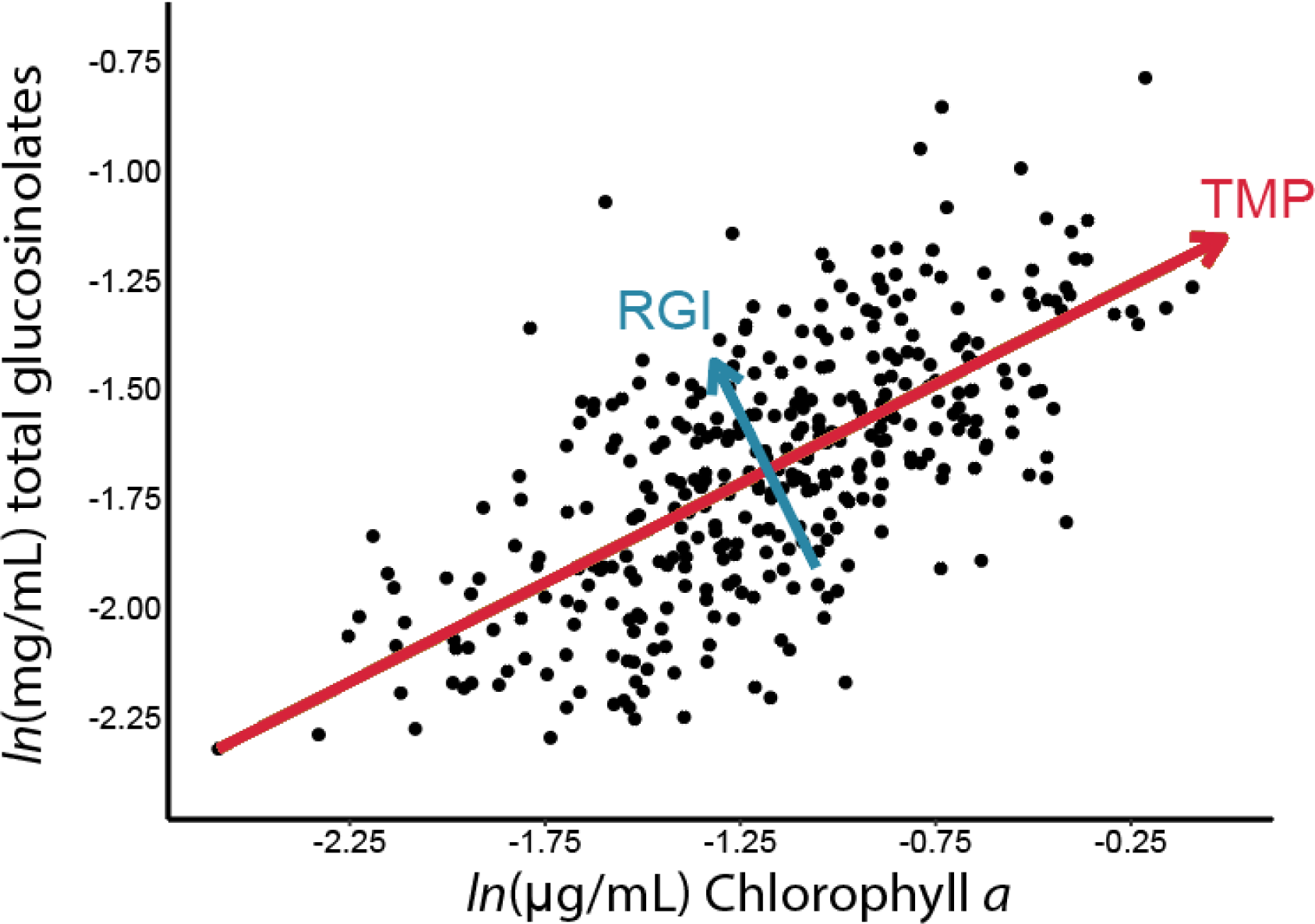
Bivariate plot of glucosinolate and Chl *a* concentrations with approximate principal component eigenvector ‘traits’ representing Total Metabolite Production (TMP, red) and Relative Glucosinolate Investment (RGI, blue). TMP accounts for 84 % of the total variation while RGI accounted for 16 % of the total variation.

### Plasticity and genetic variation

We found significant genetic variation amongst the 23 *A. petiolata* seed families for survival, transplant size (TS_AP_), lifetime fitness, glucosinolate production, and RGI, but not for TMP or Chl *a* (Table 2). Seed family explained 28.59 % of the variation in TS_AP_, 17.96 % of the variation in survival and 7.38 % of the variation in lifetime fitness (all *p* < 0.001). Notably, seed family only explained 7.47 % of variation in glucosinolate concentration, and it was not a significant predictor of TMP (*p* = 0.08) or Chl *a* production (*p* = 0.19). In contrast, the effect of seed family on RGI was highly significant (P < 0.001) and explained 16.91 % of the phenotypic variance. Of the traits with detectable genetic variation, only glucosinolate production and lifetime fitness varied significantly among treatments. Transplant size (TS_AP_) was a significant predictor of relative growth rate, total leaf area (TLA_AP_) and fecundity, but had little effect on the magnitude of variance estimates (Supporting Information Table B5). Genetic variation was not detectable in the remaining traits, which all exhibited strong plastic responses to treatment. We did not detect significant genetic variation for plasticity in any traits in the form of statistical interactions between seed family by competition treatment (Table 2).

**Table 2.** Percent of total phenotypic variation explained by family (broad sense heritability [*H^2^*]), treatment (plasticity) and a family-by-treatment interaction (genetic variation in plasticity/reaction norms).

| Variables | Family ($H^2$) | Treatment | Family × Treatment |
| --- | --- | --- | --- |
| | % ($\chi^2$, p) | % ($\chi^2$, p) | % ($\chi^2$, p) |
| Transplant size | **28.59 (107.5, < 0.001)** | 0.13 (0.3, 0.56) | 2.57 (1.4, 0.24) |
| Chl $a^{*\dagger}$ | 2.21 (1.7, 0.19) | **8.54 (27.1, < 0.001)** | < 0.1 (0, 1) |
| Glucosinolate$^{*\dagger}$ | **7.47 (14.0, < 0.001)** | **8.78 (26.3, < 0.001)** | 3.02 (0.6, 0.43) |
| Total metabolite production (TMP) † | 2.91 (3.1, 0.08) | **10.95 (32.5, < 0.001)** | 2.13 (0.3, 0.59) |
| Relative glucosinolate investment (RGI) | **16.91 (39.6, < 0.001)** | < 0.1 (0, 1) | <0.1 (0.1, 0.1) |
| Relative growth rate | 1.89 (3.2, 0.07) | **11.19 (48.9, < 0.001)** | 3.22 (1.3, 0.25) |
| TLA | 0.17 (0.9, 0.35) | **27.40 (141.0, < 0.001)** | 2.85 (1.8, 0.19) |
| Survival | **17.96 (22.3, < 0.001)** | 0.26 (0.1, 0.79) | < 0.1 (0, 1) |
| Bolt size | <0.1 (0, 1) | **14.46 (17.1, < 0.001)** | < 0.1 (0, 1) |
| Fecundity$^*$ | < 0.1 (0, 1) | **11.52 (12.0, < 0.001)** | < 0.1 (0, 1) |
| Lifetime Fitness$^*$ | **7.38 (14.2, < 0.001)** | **3.51 (11.7, < 0.001)** | 1.44 (0, 1) |

Bold indicates $p < 0.05$
$^*$ Analyzed on log scale
† Leaf area of the sampled leaf included as a fixed effect.

### Natural selection

*Alliaria petiolata* performance in year 1 (TLA_AP_) was best explained by TS_AP_ (β = +0.06, *p* < 0.001), treatment, TMP, *X. campestris* infection (β = –0.09, *p* < 0.001), fern abundance (β = – 0.05, *p* < 0.01) and the random effect of greenhouse bench (χ^2^ = 29.9, *p* < 0.001). We found a significant interaction between TMP and treatment (χ^2^ = 11.9, *p* < 0.01), indicating different slopes. Specifically, TMP was positively associated with TLA_AP_ in the interspecific treatment (β = +0.10, *p* < 0.001), but was not associated with TLA_AP_ in any other treatment (Figure 3). The interaction between TMP and treatment remained significant after controlling for leaf area of the sampled leaf (χ^2^ = 12.6, *p* < 0.01). Compared to the alone treatment, interspecific competition reduced TLA_AP_ by –15% (*p* < 0.01) while intraspecific competition reduced TLA_AP_ on average by –54 % (*p* < 0.001).

**Figure 3.**
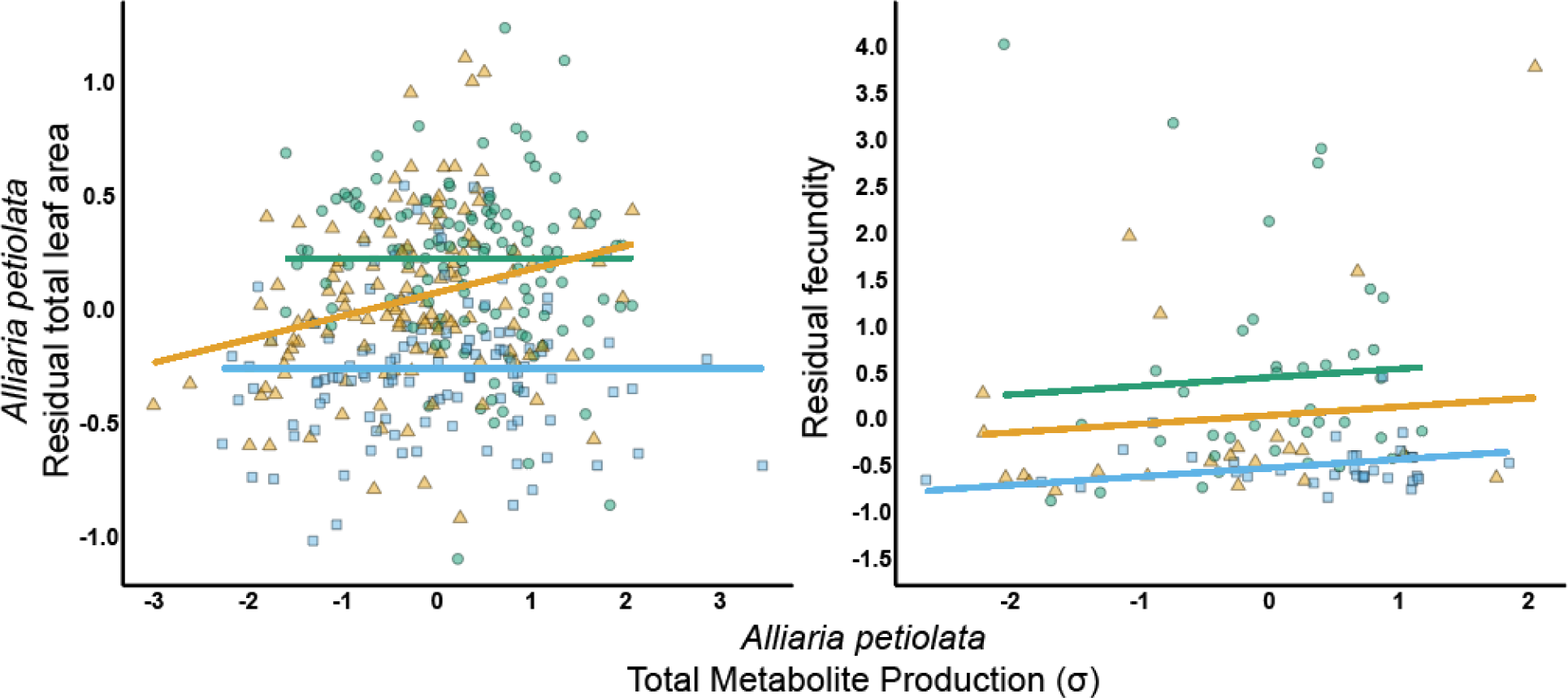
Bivariate plots of Total Metabolite Production (TMP) for *A. petiolata* against (A) total leaf area or (B) mean-standardized fecundity – a measure of relative fitness. Green circles denote the alone treatment, orange triangles denote the interspecific treatment and blue squares denote the intraspecific treatment. The y-axis in panel A shows mean-standardized total leaf area (unitless), after controlling for *X. campestris* infection, fern abundance, greenhouse bench and *A. petiolata* transplant size. The y-axis in panel B shows mean-standardized fecundity after controlling for *A. petiolata* transplant size, showing selection gradients. Both x-axes show z-standardized *A. petiolata* TMP, representing the production of both glucosinolates and Chl *a*. Slopes of zero reflect non-significant selection gradients. Regression lines are colored according to treatment.

*Alliaria petiolata* survival was best explained by fern abundance (β = +0.35, *p* < 0.001) and the random effects of family (χ^2^ = 12.3, *p* < 0.001) and field row (χ^2^ = 6.7, *p* < 0.05). The TMP by treatment interaction was significant (χ^2^ = 6.6, *p* < 0.05), such that TMP was positively associated with survival in the intraspecific treatment only (β = +0.49, p < 0.05). However, after accounting for leaf area of the sampled leaf, the effect was no longer significant (χ^2^ = 5.3, *p* = 0.07). When family was removed from the model, the effect of RGI on survival was on the verge of significance (β = –0.14, *p* = 0.059). Interspecific competition reduced *A. petiolata* survival by 33% (χ^2^ = 5.3, *p* < 0.05). In contrast intraspecific competition was associated with a 7% reduction in *A. petiolata* survival, but was not significant (*p* = 0.67).

Selection differentials of survival were obtained by determining the mean change in trait value between surviving individuals (after purging selection) and all individuals (before purging selection), in units of standard deviation. After selection, mean TMP increased by +0.10 σ and +0.43 σ in the inter- and intra-specific treatments, respectively, but remained relatively constant in the *A. petiolata* alone treatment (–0.03 σ). In contrast, RGI decreased by –0.22 σ and –0.32 σ in the inter- and intra-specific treatments, respectively, and remained relatively constant in the *A. petiolata* alone treatment (+0.01 σ).

Fecundity was best explained by TS_AP_ (β = +0.24, *p* < 0.001), TMP, and treatment (χ^2^ = 9.4, *p* < 0.01). Unlike in models of TLA_AP_ and survival, TMP was positively associated with fecundity in all treatments (β = +0.14, *p* < 0.05) (Figure 3). Compared to the alone treatment, interspecific competition reduced fecundity by 50% (*p* < 0.01), while intraspecific competition reduced fecundity by 120% (*p* < 0.001) on average. Lifetime fitness was best explained by treatment (χ^2^ = 15.5, *p* < 0.001), family (χ^2^ = 13.2, *p* < 0.001) and field row (χ^2^ = 9.0, *p* < 0.01). Interspecific competition reduced lifetime fitness by 106% (*p* < 0.001), while intraspecific competition reduced lifetime fitness by 119% (*p* < 0.001; see Table B7 for all *A. petiolata* model selection results).

### Acer saccharum *performance*

When assessing treatment effects, TLA_AS_ in year 1 was best predicted by TS_AS_ (β = +0.39, *p* < 0.001), fern abundance (β = –0.07, *p* < 0.05), *A. saccharum* leaf damage (β = –0.13, p < 0.001) and treatment (χ^2^ = 17.5, *p* < 0.001). Compared to the alone treatment, interspecific competition reduced TLA_AS_ by 12.6% on average (*p* < 0.001). In the second year, *A. saccharum* shoot mass was best explained by treatment (χ^2^ = 17.6, *p* < 0.001) and TS_AS_ only (β = +0.12, *p* < 0.001). Interspecific competition reduced *A. saccharum* shoot mass in year 2 by –40.4% on average (*p* < 0.001).

When assessing which *A. petiolata* traits affected *A. saccharum*, TLA_AS_ was best explained by TS_AS_ (β = +0.40, *p* < 0.001), *A. saccharum* leaf damage (β = –0.11, *p* < 0.001) and the two *A. petiolata* traits, TMP and RGI. While TMP was negatively associated with TLA_AS_ (β = –0.11, *p* < 0.001), RGI was positively associated with TLA_AS_ (β = +0.07, p < 0.05) (Figure 4). Shoot mass in year two was best explained by TS_AS_ (β = +0.18, *p* < 0.001), field row (χ^2^ = 5.5, *p* < 0.05), and the competitor trait, TLA_AP_ (β = –0.13, *p* < 0.001) (see Table B6 for all *A. saccharum* model selection results).

**Figure 4.**
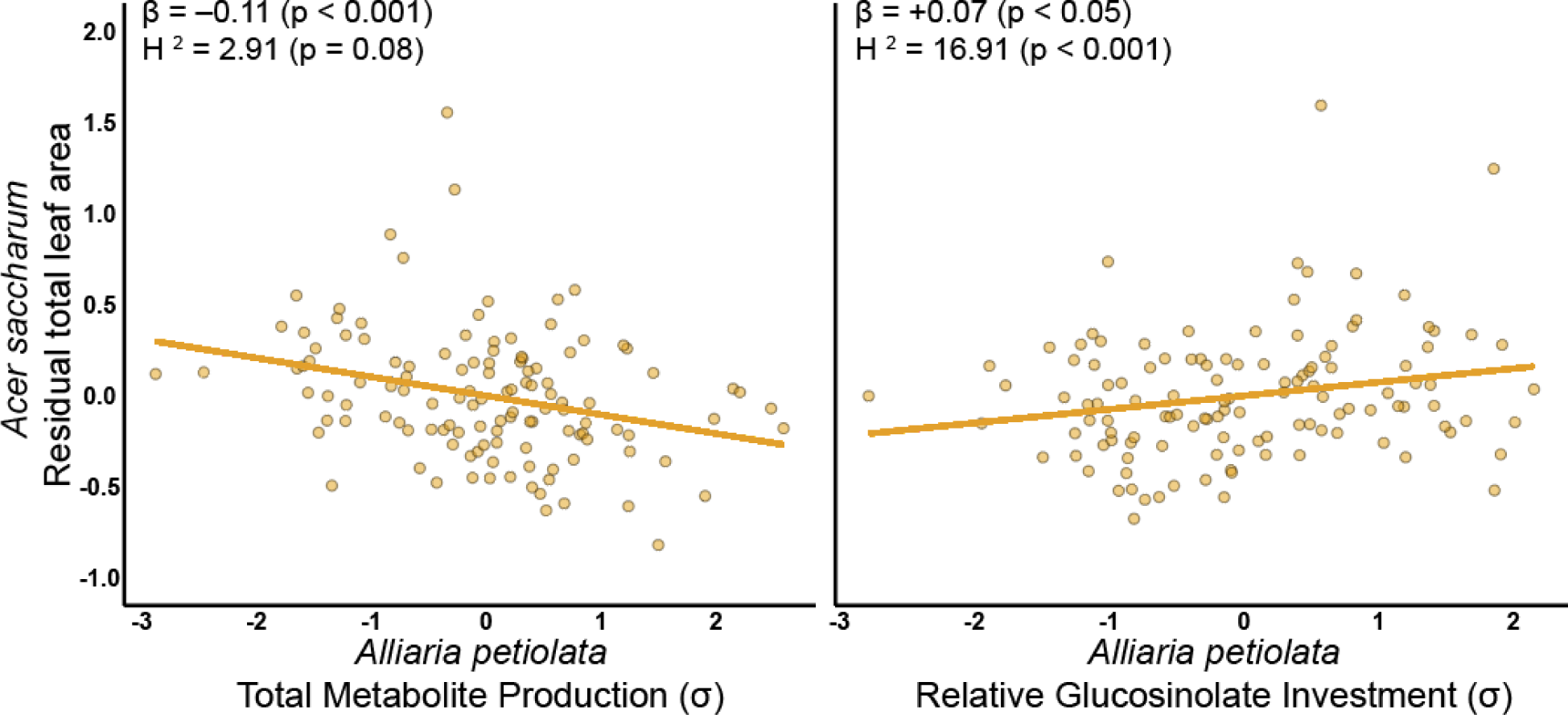
Bivariate plots of (A) Total Metabolite Production (TMP) and (B) Relative Glucosinolate Investment (RGI) against *Acer saccharum* total leaf area at the end of year 1. The y-axis shows the mean-standardized total leaf area (unitless) after controlling for *A. saccharum* transplant size and leaf damage. Both TMP and RGI were z-standardized to a mean of zero and unit standard deviation. Regression coefficients (*ß*) and broad sense heritabilities (*H^2^*) are also shown for each graph.

### Path analysis

TMP and RGI were significantly associated with TLA_AS_ in year 1, but not *A. saccharum* shoot mass in year 2. Therefore, TLA_AS_ was included in all path analyses as the intermediate step linking TMP and RGI to *A. petiolata* fitness. Significant predictors (as described in the results above) of *A. petiolata* TLA_AP_, survival, fecundity and lifetime fitness were included as covariates in path analyses of *A. petiolata* TLA_AP_, survival, fecundity and lifetime fitness, respectively. Significant predictors of *A. saccharum* TLA_AS_ were included in all path analyses.

In the TLA_AP_ path analysis, TLA_AS_ was negatively associated with TLA_AP_ (β = –0.11, *p* < 0.05). The indirect effects of TMP (β = +0.02) and RGI (β = –0.01), through TLA_AS_, on TLA_AP_ were both significant (both *p* < 0.01); however, the direct effects of TMP and RGI on TLA_AP_ were not (*p* = 0.08 and 0.19, respectively; Supporting Information Figure B1). Likewise, in the survival path analysis, TLA_AS_ was also negatively associated with *A. petiolata* survival (β = –0.38, *p* < 0.05) and the indirect effects of TMP (β = +0.07, *p* < 0.05) and RGI (β = –0.05, *p* < 0.01), through TLA_AS_, on survival were significant (Figure 5). Like in the TLA_AP_ path analysis, the direct effect of both TMP and RGI on survival were not significant (*p* = 0.83 and 0.30, respectively). Both path analyses suggest that TMP and RGI had opposite effects on *A. petiolata* fitness and only impacted fitness through their intermediate effects on *A. saccharum*.

**Figure 5.**
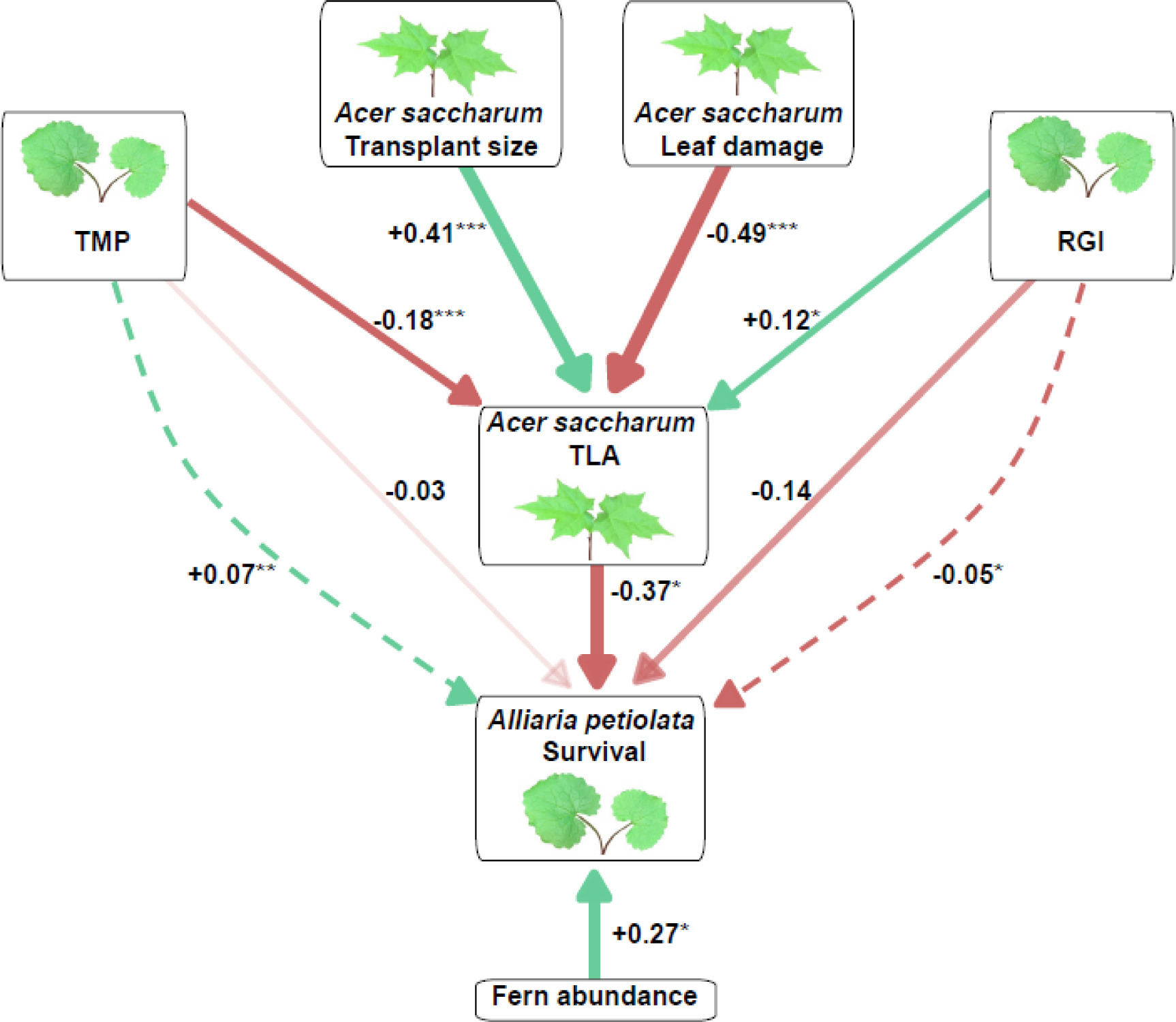
Path analysis of *Alliaria petiolata* survival. Survival was mean-standardized while other variables were z-standardized such that coefficients are at the same scale as selection gradients. Arrows denote the direction of the effect, with green arrows indicating a positive gradient and red arrows indicating a negative gradient. Arrow thickness denotes the magnitude of the effect and darker arrows indicate greater significance. The text next to each arrow shows the path coefficient and its significance level (*p* < 0.05*, *p* < 0.01**, *p* < 0.001***). Total Metabolite Production (TMP) denotes the production of both glucosinolates and Chl *a*, while Relative Glucosinolate Investment (RGI) indicates the production of glucosinolates relative to Chl *a*. *Acer saccharum* TLA denotes the total leaf area of *Acer saccharum* at the end of year 1. Solid arrows represent direct effects while dashed arrows represent indirect effects.

In contrast, TLA_AS_ was not associated with fecundity (*p* = 0.92) and there were no significant direct or indirect effects of TMP or RGI on fecundity (Supporting Information Figure B2). Similarly, TLA_AS_ was not a significant predictor in the path analysis of lifetime fitness (*p* = 0.28), and there were no indirect effects of TMP or RGI on *A. petiolata* lifetime fitness. There was a significant direct effect of RGI on lifetime fitness (β = –0.50, *p* < 0.05), but not of TMP (*p* = 0.08; Figure 6).

**Figure 6.**
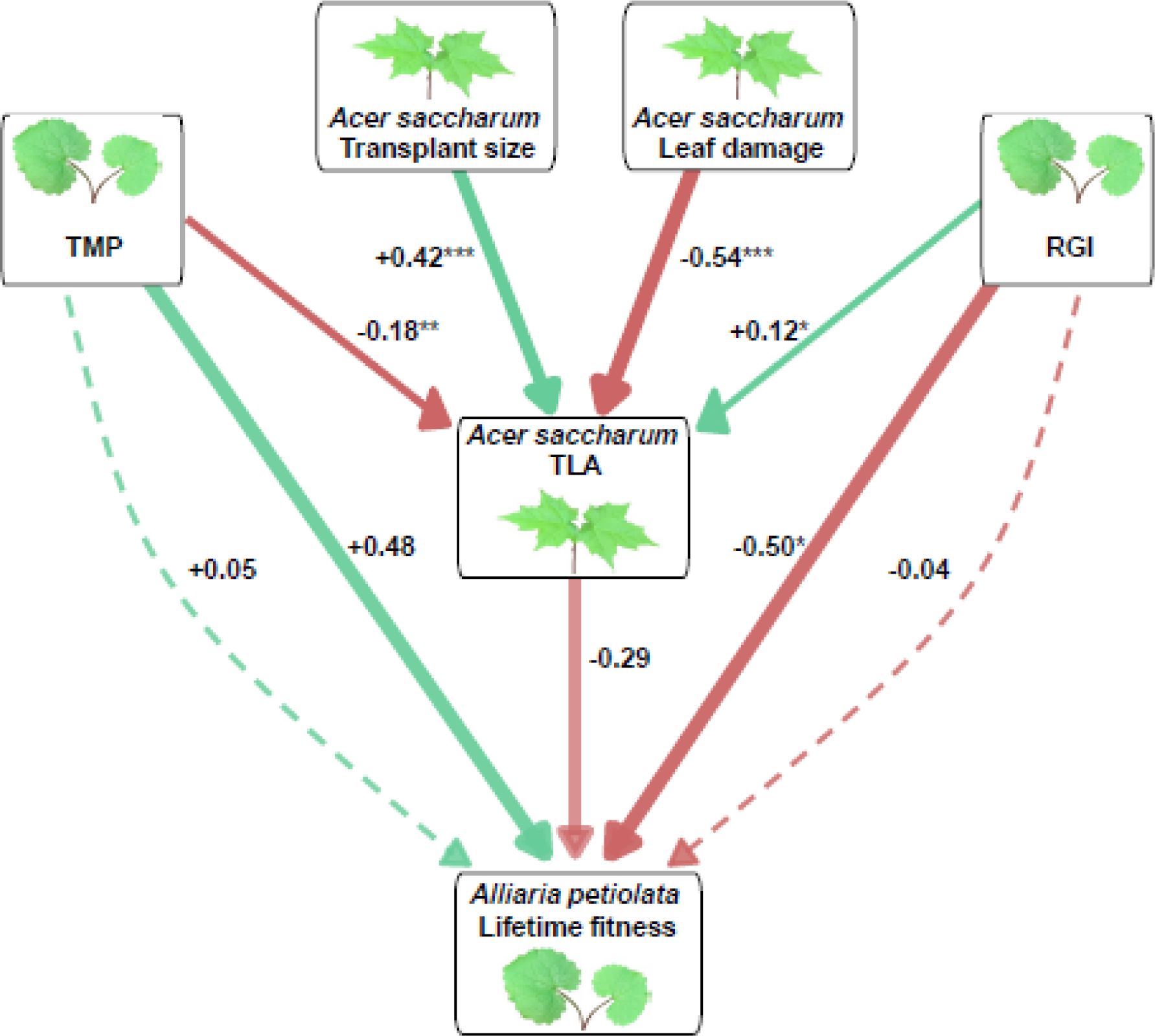
Path analysis of *Alliaria petiolata* lifetime fitness. Scaling and aesthetics are the same as described in Figure 5.

## Discussion

Plants that invade and spread across large geographical areas experience heterogeneous biotic and abiotic environments (van Kleunen et al. 2018). In the Brassicaceae family, glucosinolates play important roles in plant responses to biotic interactions and abiotic stress (Holst and Fenwick 2003). To investigate glucosinolate evolution and the potential for eco-evolutionary dynamics in glucosinolate production via allelopathy, we quantified plasticity, heritability, and natural selection via direct and indirect ecological effects across three competition treatments.

Strong trait correlations compromise estimates of selection gradients, which can be addressed using principal components analysis (Chong et al. 2018). As predicted, we found a strong, positive correlation between glucosinolate concentration and Chl *a* (Figure 2). A PCA resulted in two statistically independent axes that fortuitously differed in heritability. The first principal component corresponded to plastic variation in Total Metabolite Production (TMP), while the second delineated heritable variation in Relative Glucosinolate Investment (RGI) (Table 2). This relatively simple analysis captured genetic variation from a native genotype from Italy and 22 populations of different age structure, density, habitat, herbivory, and pathogen damage, spanning ∼3,000 km across Canada and the United States, from British Columbia and Colorado to Georgia and Maine. Below we discuss implications of plastic variation in TMP, and heritable variation in RGI, for the evolution of allelopathy during *A. petiolata* invasion, and how these might translate to other invasive species.

The eco-evolutionary feedbacks hypothesis (Lankau et al. 2009; Evans et al. 2016) makes at least three predictions that can be evaluated with our experiment. First, glucosinolate production should be highly heritable, particularly when assessed across populations evolved under different degrees of intra- and interspecific competition. Contrary to this prediction, we found that glucosinolate concentrations were highly plastic across competition treatments, with relatively low heritability (Table 2). RGI was significantly heritable in the broad sense (*H*^2^ = 39.6), but only explained 16% of the variation in glucosinolates and Chl *a*. The second prediction is that natural selection should favour higher investment into glucosinolate production when phenotypes experience interspecific competition, and lower investment under intraspecific competition or no competition. Instead, we found that natural selection favors genotypes with lower RGI. Specifically, the selection differential for RGI was -0.22 σ and -0.32 σ in the inter- and intra-specific treatments, respectively, and there was no selection detected in the no-competition treatment (+0.01 σ). The third prediction is that glucosinolate investment should increase fitness of *A. petiolata* indirectly by suppressing growth of competing *A. saccharum*. Instead, we found a positive correlation between RGI and the growth of naïve *A. saccharum* (Figure 4), resulting in an indirect negative correlation with *A*. *petiolata* fitness (Figure 5). The predicted correlations were found for TMP, but adaptive evolution is unlikely for this trait given its low heritability. Instead, the adaptive evolutionary response to natural selection under both intra- and interspecific competition should produce populations that invest less into glucosinolate production. In summary, none of our main results support the hypothesis that competition dynamics during invasion drive eco-evolutionary feedbacks that maintain genetic variation in glucosinolate production across the introduced range.

Although we do not find evidence to support the eco-evolutionary feedbacks hypothesis, the strong selection gradients and high level of phenotypic plasticity observed in TMP are relevant to understanding invasion dynamics of *A. petiolata*. Environmental conditions and ecological interactions are likely more variable in natural populations than our relatively controlled greenhouse and field gardens, which suggests that plasticity in natural populations should be both common and ecologically important. We therefore conclude that the evolutionary potential of TMP is minimal, even though it increases *A. petiolata* fitness. Instead, TMP represents a highly plastic trait that can explain variation in performance among *A. petiolata* populations in North America.

In addition to competitive interactions, plasticity in glucosinolate production is likely affected by the presence of herbivores. This is relevant because glucosinolate production is upregulated by generalist herbivores and passed on to offspring (Holeski et al. 2012). Previous experiments using herbivore exclosures in the northeastern United States reveal competitive effects between *A. petiolata* and native *Trillium erectum*, and preferential herbivory on the latter (Kalisz et al. 2014; Bialic-Murphy et al. 2019. We found an increase in TMP competition was reduced, as would be the case in a population experiencing herbivory. As such, the apparent competitive advantage of *A*. *petiolata* in natural populations may not be caused by glucosinolate production, but rather both glucosinolate production and population density may be spuriously correlated due to shared effects of herbivory. We might call this hypothesis ‘apparent allelopathy’ as a special case of ‘apparent competition’ as defined by Holt (1977). Our study was not designed to test effects of herbivory, but future experiments should consider rearing at least one generation in a herbivore-free common garden to mitigate the effects of transgenerational plasticity on glucosinolate production.

In contrast to TMP, we found heritable genetic variation for RGI, but selection favoured a lower relative investment into glucosinolate production under competition. This finding is contrary to the eco-evolutionary feedback hypothesis but may be important for understanding the causes and consequences of glucosinolate evolution during invasion of *A*. *petiolata*. The fitness cost of RGI under competition should favor genotypes with lower investment into glucosinolates, resulting in evolution of lower RGI. The maintenance of heritable genetic variation in RGI, despite this fitness cost, is consistent with glucosinolate investment playing other important ecological roles during invasion. For example, the evolution of increased competitive ability (EICA) hypothesis predicts an evolutionary reallocation of resources from defense to growth and competition in populations with fewer natural enemies (Blossey and Notzold 1995). Indeed, *A petiolata* may have fewer natural enemies in North America compared to Europe (Rodgers et al. 2008), and total glucosinolate production in North America was estimated to be 36% lower than native European populations (Cipollini et al. 2005). However, framing this hypothesis within a native-introduced dichotomy overlooks considerable variation among introduced populations, which can be higher than the average difference between ranges (e.g., compare population vs range means in Figure 1 of Cipollini et al. 2005). If all introduced populations escaped regulation by specialist enemies, then we would not expect significant heritability of RGI across populations as we observe in this study. Instead, heritable RGI is more consistent with herbivore communities that are heterogeneous throughout eastern North America, maintaining genetic variation in glucosinolate investment. In addition to herbivory, glucosinolates are known to play other ecological roles that could help maintain genetic variation in RGI, including response to abiotic stress (Louda and Mole 1991). Testing the effects of herbivory and biotic stress is beyond the scope of our experiment, but the same seed families that we used are available from the Global Garlic Mustard Field Survey (Colautti et al. 2014) to enable such studies in the future.

In summary, we examined plasticity, heritability, and natural selection of glucosinolate production in one native and 22 introduced populations of *A. petiolata*, reared in three competition treatments in greenhouse and field common gardens. We found that most traits were highly plastic, and selection consistently favored genotypes that invest less into glucosinolate production relative to Chl *a* in competitive environments. We conclude that eco-evolutionary feedbacks involving competition have not likely played a significant role in the invasion of *A. petiolata* across eastern North America. Instead, the successful spread of this species more likely owes to preadapted traits and phenotypic plasticity. Although not involved in competition, glucosinolates likely play other ecological roles during invasion that can be tested with seed accessions available from the Global Garlic Mustard Field Survey. More generally, our results demonstrate how evolution during biological invasion can be better understood through the application of quantitative genetics, selection gradients, and causal analysis.

## Author Contributions

RH and RIC conceived and designed the experiment. RH and MM ran the experiment with financial support and research assistants provided by RIC. RH led the data analysis manuscript writing with contributions from MM and writing guidance from RIC.

## Acknowledgements

The authors are grateful to M. Fuentes, D. Felstead, L. Wisteard, J. Stollman, C. Smith, and E. Newman for helping with experiment set up, sample processing and data collection. We are also grateful to Drs. F. Bonier and L. Aarssen for helpful suggestions for the thesis on which this manuscript is based. Additional critiques and suggestions from Y. Buckley and two anonymous reviewers significantly improved the manuscript. Funding was provided to RIC from NSERC Discovery and from a CFI Tier II Canada Research Chair.

## Data Availability

All original data and reproducible code are available through Dryad (https://doi.org/10.5061/dryad.cfxpnvxbj).

