## Supporting Information for "Direct and indirect fitness effects of plant metabolites and genetic constraints limit evolution of allelopathy in an invading plant"

### Supporting Information for Honor, Marcellus and Colautti "Direct and indirect fitness effects of competition limit evolution of allelopathy in an invading plant"

For raw data and detailed code, see Dryad: https://doi.org/10.5061/dryad.cfxpnvxbj

#### Appendix A (Supplementary Methods)

***Study species***

The biennial herb *Alliaria petiolata* (garlic mustard) is emerging as a model system for integrated studies in invasion biology and ecological genetics more generally (Kueffer et al. 2013; Colautti et al. 2014; Alabi et al. 2021). A member of the mustard family (Brassicaceae), *A. petiolata* is native to much of Eurasia and invasive in much of North America, where it can be an aggressive weed (Rodgers et al. 2008). As an obligate biennial, it emerges in early spring and overwinters as a rosette before producing a flowering bolt in the following spring (Cavers et al. 1979; Anderson et al. 1996; Cruden et al. 1996). *Alliaria petiolata* reproduces sexually but over 80% of seeds are produced through self-fertilization and roughly 80% of the genetic variation for neutral markers is partitioned among populations (Durka et al. 2005). Despite high selfing rates, the species does not tend to suffer from inbreeding depression, (Mullarkey et al. 2013) possibly due to its hexaploid genome, genetic purging, and periodic outcrossing. Members of the Brassicaceae family produce glucosinolates for a variety of ecological purposes including defence against pathogens and herbivores, and tolerance to other environmental stresses (Holst and Fenwick 2003; Hopkins and Hüner 2009). Glucosinolates produced by *A. petiolata* enter the soil through shoot decomposition and root exudation (Barto and Cipollini 2009; Cantor et al.

2011), which can disrupt mycorrhizal communities and reduce mycorrhizal colonization of host plants (Roberts and Anderson 2001; Stinson et al. 2006; Callaway et al. 2008; Wolfe et al. 2008; Cantor et al. 2011). However, these allelopathic effects are inconsistently observed, depending on experimental conditions and the presence of susceptible neighbours and mycorrhizae (Burke 2008; Barto et al. 2011; Burke et al. 2019; Duchesneau et al. 2021; Roche et al. 2021). Understanding the context dependency of glucosinolate-soil interactions remains an active area of research in this species.

*Acer saccharum* (sugar maple) is a shade tolerant tree that is native to over 25 different forest types in North America; it typically grows in cool moist regions between the latitudes of approximately 35° and 50° (Ralph 1998). Its seeds exhibit very high germination success (~95%) and densities regularly exceed 37 saplings per $m^2$ in the forest understory (Godman et al. 1990). The high sapling density and reliance on root mycorrhizal associations make saplings of *A. saccharum* a primary competitor of *A. petiolata* and a good model for allelopathy studies (Stinson et al. 2006). Early molecular work identified reduced root colonization by mycorrhizae where the two species co-occur (Barto et al. 2011), though detailed high-throughput sequencing studies have not detected an effect of *A. petiolata* on soil endomycorrhizal communities (Duchesneau et al. 2021; Edwards et al. 2022).

**Experimental setup**

*Alliaria petiolata* seeds were stratified in the dark at 4°C in Parafilm™-sealed petri dishes containing 2mL of deionized water and untreated field soil, to maintain characteristics and microbiota of natural soil. We monitored germination from March 7–May 7, 2019. Seeds that did not germinate by March 26, 2019, were forced to germinate using scarification and 2mL of 0.001M gibberellic acid, as per Sosnoskie and Cardina (2009).

Upon germination, we transferred seedlings into 2cm x 2cm plug trays filled with a 1:1 mixture of sand and Sun Grow Sunshine® Mix #2 potting soil, which does not contain added nutrients. Trays were watered as needed with deionized water to maintain soil saturation. To limit variation in growth among seed families with different germination times, we maintained seedlings at a constant 10°C (12h half daylight) in a growth chamber in the Queen's University Phytotron.

We transplanted all available seedlings on May 9–10, 2019, into 4-inch plastic pots (surface area = 81cm$^2$; volume = 524.39 cm$^3$) containing a 1:1 mixture of field soil and Sun Grow Sunshine® Mix #2 potting soil. Pots were moved to a glasshouse room in the Queen's University Phytotron, and we randomized the location of each pot across 5 tables. In the glasshouse, plants received natural light, filtered with a shade cloth to simulate a forest overstory, and were broadcast watered with a rubber hose once or twice per day as needed to maintain soil that was moist to the touch.

To determine the competitor pairs to be used in the intraspecific treatment, we wrote a short algorithm in R to randomly pair individuals from different seed families without repeating pair combinations.

To characterize transplant size (TS) for *A. saccharum* ($TS_{AS}$) and *A. petiolata* ($TS_{AP}$), we used principal components analysis (PCA). The $TS_{AS}$ PCA included stem height, largest leaf length and number of leaves, all measured after transplant (May 16 and 17, 2019), as well as stem height, number of leaves and $TLA_{AS}$, all measured at the end of year 1 (August 14 to 21, 2019), (Table B1). The $TS_{AP}$ PCA included initial largest leaf length and number of true leaves, measured at transplant (Table B2). The first principal component from both PCAs were used to represent the size of both plants at transplant.

**Machine learning analysis of plant size**

To measure rosette size in *Alliaria petiolata*, we trained a machine learning model of whole plant images. The top and side of each *A. petiolata* rosette were photographed using two webcams affixed to a custom image-capture stand (Figure A4). Cameras were oriented to image the top and side of each rosette to capture leaf area in all three dimensions. We measured relative plant size as the total pixel area of each plant, given that cameras were fixed at a constant distance from each imaged plant, (Figure A5).

Image analysis was performed using a custom machine learning algorithm written in Python 3.8 using the packages OpenCV2 and PlantCV2 (Bradski 2000; Python Core Team 2015; Berry et al. 2018). We first trained the computer to identify leaves based on the color spectrum of each pixel captured by the camera, encoded on an rgb scale (0-255). In a subset of 100 training images, we manually assigned 10 pixels in each image as leaves or background. From this training set, a Naïve Bayesian classifier algorithm available in the PlantCV2 package could assign a posterior probability of leaf vs background to all pixels in the remaining images.

A subset of 76 plants were used to verify the machine learning estimates. For these plants, total leaf area (TLA) was estimated manually by measuring the area of each leaf (defined as leaf length times width) and summing across all leaves for each plant. There was a strong correlation between rosette size (measured as the total number of pixels in an image corresponding to plant) and TLA ($p < 0.001$, $R^2$=0.852), which allowed us to automate measurements of all 5,659 photos. Each photo that was analyzed produced an output image, outlining the parts identified as rosette (i.e., Figure A5). Each of the 5,659 images were manually inspected to check for assignment errors in the machine learning algorithm. We found that 29 plants were mis-identified. In these instances, we sampled pixels directly from the mis-identified images and retrained the machine learning algorithm on these pixels until all plants were correctly identified.

We wrote a program in Python 3.8 to automate image capture. This program displays webcam images to assist with correct placement of each plant, allows the user to take a photo with each camera, and displays the photos for manual quality inspection. If approved by the user, the images are automatically stored and named with the appropriate identification (as read through a barcode scanner) and the exact time that the photos were taken. We used Unix time, which is the number of seconds since Jan 1st, 1970, providing a precise time of measurement with which we could calculate growth rates with high accuracy.

Plants were imaged every two weeks over the course of two days, but most growth occurred between the first (May 27 and 28) and second (June 11 and 12) censuses. We calculated relative growth rate between these two time periods and used this variable in analyses:

$$\text{Relative growth rate} = \frac{\ln(Rosette\ size_2) - \ln(Rosette\ size_1)}{t_2 - t_1} \qquad 1$$

where $t$ denotes the time in seconds (Unix time), and subscripts indicate the census.

**Secondary compound processing and analysis**

To quantify glucosinolate and Chl *a* concentration in *A. petiolata*, we sampled leaf tissue at the end of the first growing season (August 14 to 21, 2019). Two leaf punches of 8 mm diameter from a single leaf were immediately placed in an Eppendorf® 1.5mL flip-cap tube and flash-frozen in liquid $N_2$ prior to storing at -80 °C. Up to two leaves were sampled per plant, and the two smallest leaves that were greater than 30 mm in length were chosen for sampling from each plant. We excluded leaves with more than 50% senescence and/or pathogen damage. Leaf area (length × width) was also recorded for each leaf sampled, hereafter referred to as leaf area of the sampled leaf, to account for dilution effects associated with leaf cell expansion (Cipollini and Gruner 2007).

Leaf tissue was kept frozen and pulverized using two cycles of 15 seconds using a Next Advance Bullet Blender® at maximum speed (6 kHz). We submerged in liquid $N_2$ between cycles to limit thawing of the plant tissue. After pulverization, we added 0.55 mL of 100 % methanol and incubated in an Eppendorf Mixmate Vortex shaker for 1 hour at 25 °C and 300 rpm, followed by centrifugation for 2 minutes at 2500 g (Eppendorf, 5427 R). Next, the pellet was discarded and 485µL of supernatant was stored at –80°C until further processing.

Once ready for more processing, Chl *a* was measured by transferring 20µL of supernatant into the wells of a 96 well UV flat bottom plate; the sample was then evaporated and resuspended in

50µL of 100% methanol immediately prior to reading. Chl *a* concentration was quantified with UV spectroscopy following Ritchie (2006):

$$\text{Chl } a\left(\frac{\mu\text{g}}{\text{ml}}\right) = -8.0962(A_{652} - A_{750}) + 16.5169(A_{665} - A_{750}) \quad 2$$

where *A* is the absorbance and subscripts denote the wavelength in nanometers (nm).

Glucosinolate concentration was measured by rinsing 465 µL of supernatant with 650 µL of hexane to remove hydrophobic pigments that overlap with the target wavelength (Haribal and Renwick 2001; Callaway et al. 2008). To rinse, samples were placed on an incubator at 30°C for 25 minutes at 600rpm. Samples were then chilled in an ice bath for 15 minutes and subsequently centrifuged at 2500g for 10 minutes at 4°C.

Glucosinolate concentration of the rinsed supernatant was quantified by transferring 100µL of the sample into the wells of a 96 well UV flat bottom plate. Samples were left to evaporate and, once dry, were resuspended in 100µL of 100% methanol immediately prior to reading. The assays were read at 425nm before and after adding 100µL of 3.5 mM sodium tetrachloropalladate ($NaCl_4Pd$) to wells. This procedure followed methods in Ishida et al. (2011) and Ishida et al. (2012), which we modified by using 100 µL of 3.5 mM sodium tetrachloropalladate ($NaCl_4Pd$) instead of 3 mL of 2 mM $NaCl_4Pd$, to enable assays in a 96-well plate (Mawlong et al. 2017). Two technical replicates were measured for each composite sample of leaf tissue.

Known concentrations (0.0625, 0.1250, 0.2500, 0.5000, and 1.0000 mg/ml) of sinigrin were used as a standard as it is one of the most abundant glucosinolates in *A. petiolata* and is suspected to have allelopathic effects (Vaughn and Berhow 1999). Because assays were read before and after adding volume of $NaCl_4Pd$ (where the volume of the sample before adding $NaCl_4Pd$ was 100 µL and the volume after was 200 µL), substrates in the first reading were twice as concentrated. Therefore, we divided the first reading by 2 and subtracted this reading from the second to estimate total glucosinolate concentration:

$$\text{Total glucosinolate (mg/ml)} = 0.69\left(A_2 - \frac{A_1}{2}\right) - 0.15 \qquad 3$$

Where $A$ is the absorbance at 425 nm and subscripts denote before ($A_1$) and after $NaCl_4Pd$ was added ($A_2$). Glucosinolate concentration was estimated from a standard curve of sinigrin (Figure A3).

As a quality control, we generated a large sample of *A. petiolata* leaf tissue extract (in 100% methanol) to be processed and used as a standard sample in every 96-well assay we performed. If the glucosinolate concentration of the standard sample in any 96-well plate assays was more than 2.5 standard deviations from the mean glucosinolate concentration of the standard sample, the entire 96-well plate assay was discarded. Some samples were also lost due to cracking in the flip-cap tube during processing, or were excluded from sampling because they had observable pathogen damage or leaf senescence. Of the 506 plants in the experiment, 267 plants had samples from two leaves, 98 plants had samples from one leaf, and 141 plants had no samples.

Glucosinolate and Chl *a* concentrations were averaged in plants with samples from two leaves for analyses.

We focused on total shoot glucosinolates as a high-throughput and non-destructive measure of glucosinolate production, allowing us to also measure lifetime fitness for the same individuals. While allelopathy may be mediated through root exudates, our approach is consistent with previous studies demonstrating allelopathy in *A. petiolata* using either leaf or whole plant extracts as glucosinolate concentrations are correlated among tissues of the same plant (McCarthy and Hanson 1998; Roberts and Anderson 2001; Stinson et al. 2006; Callaway et al. 2008). While sinigrin is secreted through roots and proposed to be the causal agent of allelopathy in *A. petiolata* (Cantor et al. 2011; Evans et al. 2016), it is also the most abundant glucosinolate in *A. petiolata* (Vaughn and Berhow 1999; Haribal and Renwick 2001) and therefore should be correlated with our estimates of total glucosinolates.

**Data Analysis**

Fecundity, lifetime fitness, fern abundance, thrip damage, *X. campestris* infection, glucosinolate concentration, and Chl *a* concentration were each log-transformed to meet assumptions of normality. If total metabolite production (TMP) or relative glucosinolate investement (RGI; please see manuscript for a description of these terms) was a significant predictor, the model was reassessed after controlling for leaf area of the sampled leaf, because plant metabolites are most concentrated in younger, smaller leaves, making TMP and RGI potentially autocorrelated with performance and fitness (Cipollini and Gruner 2007). When family was a significant predictor, a separate model excluding family was used when we wanted to include effects of heritable genetic variation.

As noted in the main methods, we used PCA to identify TMP and RGI, and used these variables in a path analyses, which were assessed using permutation tests in R. Each permutation test involved resampling either TMP, RGI or *A. saccharum* performance, refitting the model with these data, extracting the test statistic, z, and taking its absolute value to generate a sampling distribution of z. The proportion of random z-values greater than the true z-value gave the p-value ($\alpha = 0.05$). Separate tests were performed for the effect of *A. saccharum* performance on *A. petiolata* fitness, the direct effects of TMP and RGI on *A. petiolata* fitness, and the indirect effects of TMP and RGI on *A. petiolata* fitness through *A. saccharum* performance.

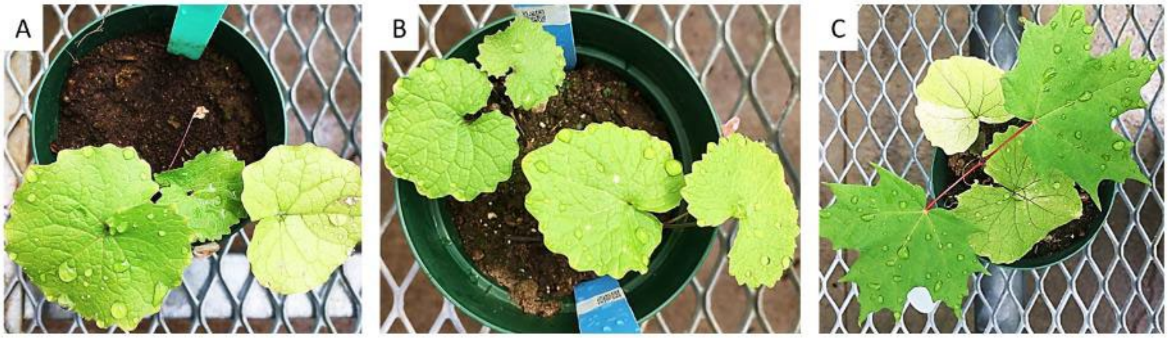


**Figure A1.** *Alliaria petiolata* grown in the (A) no competition treatment (B) intraspecific competition treatment and (c) interspecific competition treatment.

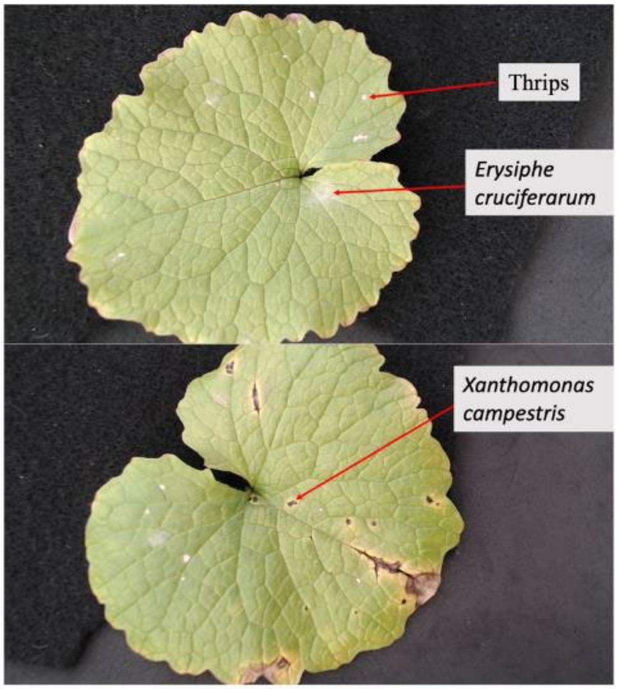


**Figure A2.** Examples of leaf images used to measure pathogen data.

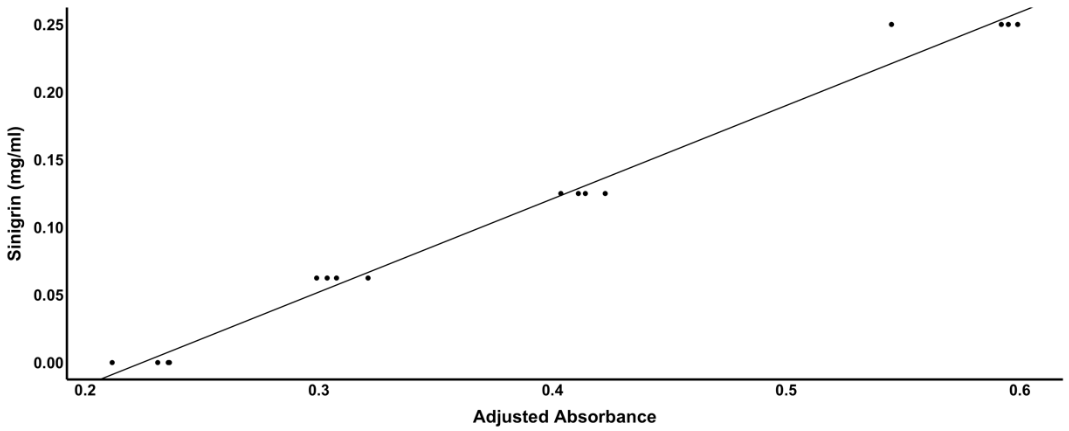


**Figure A3.** Standard curve of sinigrin concentration. $\mathbf{Sinigrin\ (mg/ml)} = \mathbf{0.69}\left(\boldsymbol{A_2} - \frac{\boldsymbol{A_1}}{\mathbf{2}}\right) - \mathbf{0.15}$, Where $\boldsymbol{A}$ is the absorbance at 425 nm and subscripts denote before ($\boldsymbol{A_1}$) and after $NaCl_4Pd$ was added ($\boldsymbol{A_2}$).

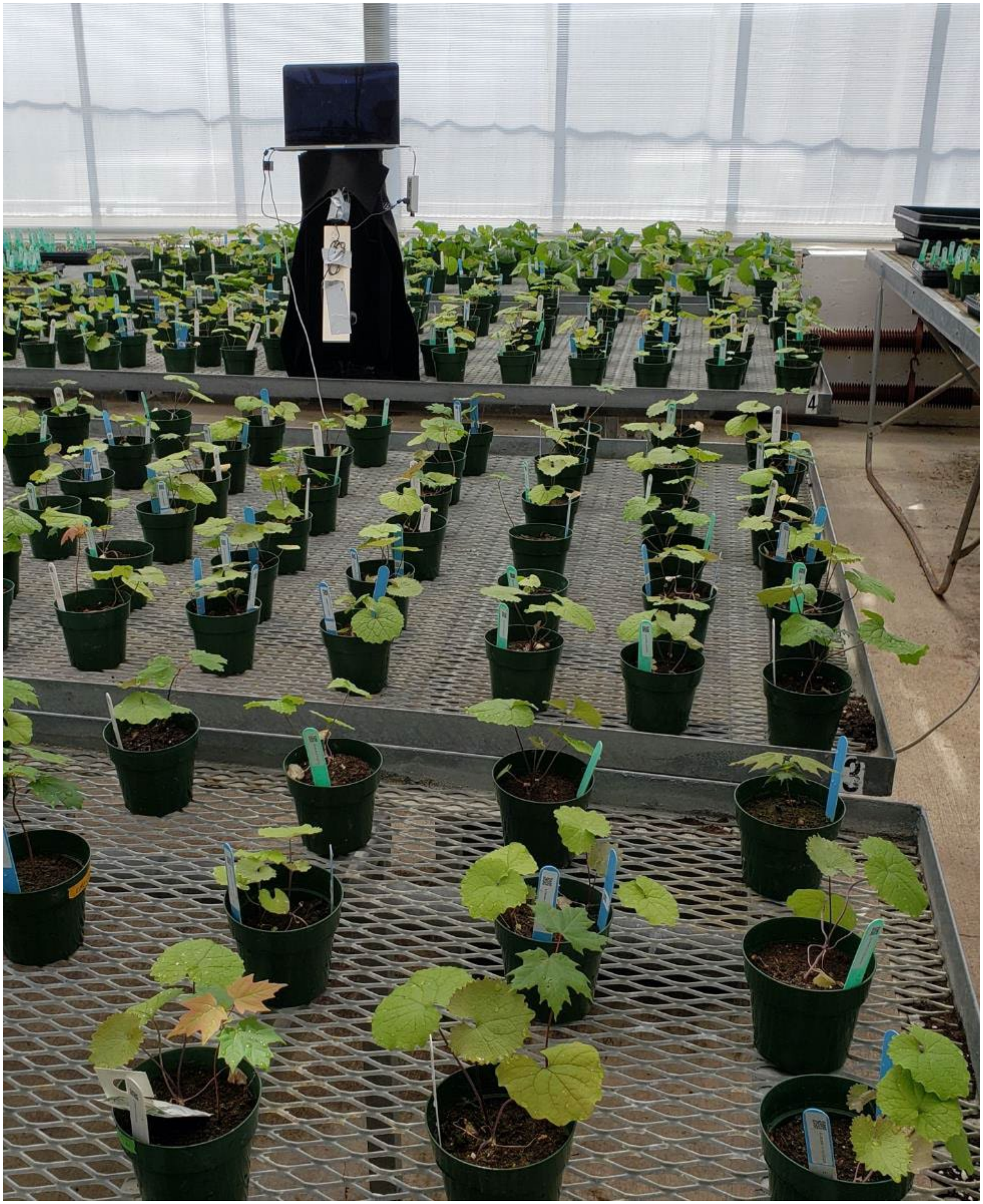

**Figure A4.** Example of greenhouse and photo apparatus used to capture images for growth rate analysis.

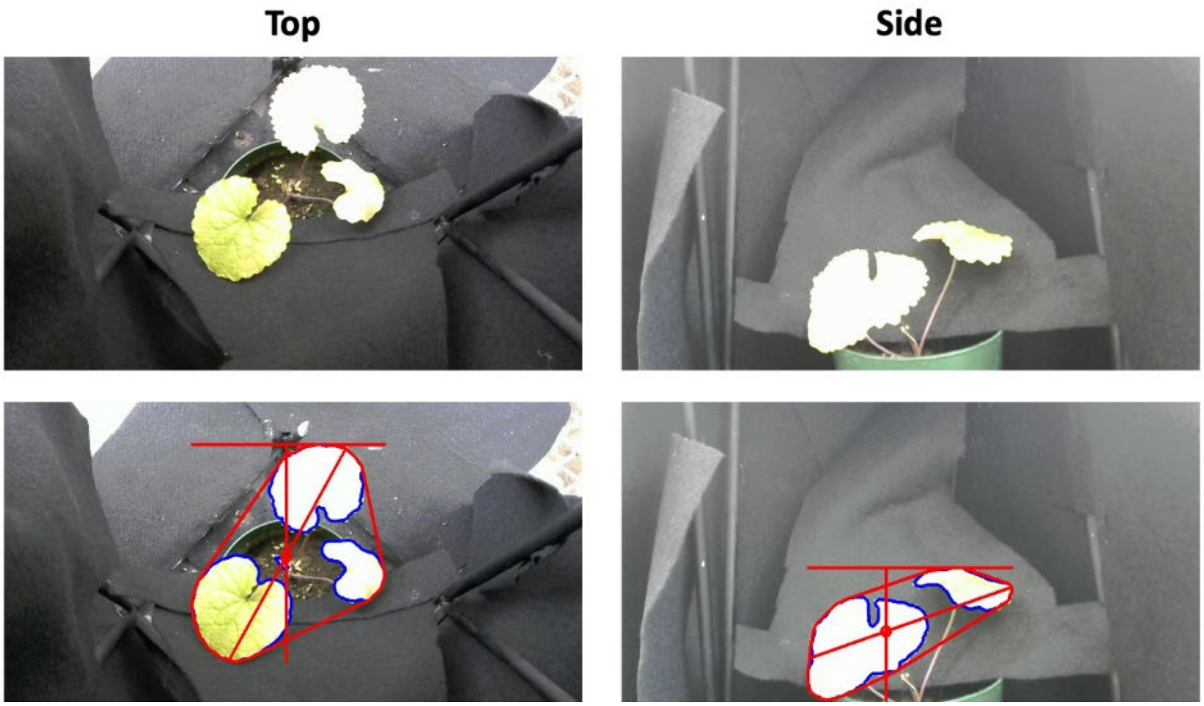


**Figure A5.** Example of plant identification with machine learning used to determine growth rate.

### Appendix B (Supporting Information)

**Table B1.** Principal component loadings of *Acer saccharum* transplant size.

| Principal Component | Loadings | | | | | | Proportion of variance |
|---|---|---|---|---|---|---|---|
| | Stem Height (initial) | Largest leaf length (initial) | Number of Leaves (initial) | Stem Height (year 1) | Number of Leaves (year 1) | Total leaf area (year 1) | |
| PC1 (Transplant size) | 0.45 | 0.40 | 0.26 | 0.53 | 0.25 | 0.48 | 0.41 |
| PC2 | 0.42 | 0.09 | -0.02 | 0.32 | -0.7 | -0.47 | 0.23 |
| PC3 | -0.18 | 0.62 | -0.68 | -0.1 | -0.23 | 0.26 | 0.17 |
| PC4 | 0.37 | -0.49 | -0.69 | 0.26 | 0.31 | -0.02 | 0.13 |
| PC5 | 0.65 | 0.18 | 0.17 | -0.72 | 0.14 | -0.07 | 0.04 |
| PC6 | -0.11 | 0.43 | -0.04 | 0.18 | 0.54 | 0.69 | 0.02 |

**Table B2.** Principal component loadings of *Alliaria petiolata* transplant size.

| **Principal Component** | **Loadings** | | **Proportion of variance** |
|---|---|---|---|
| | **Largest Leaf Length** | **Number of Leaves** | |
| PC1 (Transplant size) | 0.71 | 0.71 | 0.74 |
| PC2 | 0.71 | –0.71 | 0.26 |

**Table B3.** Summary statistics of *Alliaria petiolata* characteristics for each of the three competition treatments.

| Characteristic | Alone | Interspecific | Intraspecific |
|---|---|---|---|
| Greenhouse mortality | 1.76% (3/170) | 1.20% (2/167) | 3.74% (7/187) |
| Field mortality | 65.27% (109/167) | 73.62% (120/163) | 68.18% (120/176) |
| Relative growth rate | $1.07\times10^{-6} \pm 0.06\times10^{-6}$ | 1.01 ± 0.07 | 0.70 ± 0.05 |
| TLA ($cm^2$) | 9563 ± 474 | 8547 ± 499 | 5120 ± 384 |
| Fecundity (g) | 3.13 ± 0.84 | 1.78 ± 0.83 | 1.07 ± 0.38 |
| Glucosinolate (mg/mL) | 0.22 ± 0.01 | 0.18 ± 0.01 | 0.20 ± 0.01 |
| Chl *a* (µg/mL) | 0.39 ± 0.02 | 0.30 ± 0.02 | 0.34 ± 0.03 |
| PC1 (total metabolite production) | 0.47 ± 0.19 | –0.43 ± 0.21 | –0.05 ± 0.26 |
| PC2 (relative glucosinolate production) | 0.01 ± 0.10 | –0.01± 0.10 | 0.00 ± 0.11 |
| Leaf area of sampled leaf ($cm^2$) | 3235 ± 167 | 3194 ± 160 | 2141 ± 123 |
| *Xanthomonas campestris* (N) | 1.60 ± 0.55 | 1.08 ± 0.41 | 1.64 ± 0.66 |
| Thysanoptera (N) | 2.87 ± 0.61 | 2.82 ± 0.63 | 4.38 ± 1.49 |
| *Erysiphe cruciferarum* | 35.10% (53/151) | 20.80% (31/149) | 26.80% (41/153) |
| Fern(N) | 0.56 ± 0.33 | 1.18 ± 0.51 | 0.65 ± 0.50 |
| Lifetime fitness | 1.03 ± 0.35 | 0.40 ± 0.21 | 0.32 ± 0.13 |

Data in brackets denote the fraction used to calculate the percent. Plus or minus (±) values denote Wald's 95% confident interval (CI) around the mean in each treatment.

**Table B4.** Principal component loadings of *Alliaria petiolata* total metabolite production and relative glucosinolate investment.

| **Principal Component** | **Loadings** | | **Proportion of variance** |
|---|---|---|---|
| | **Chl *a*** | **Glucosinolate** | |
| PC1 (Total metabolite production) | 0.71 | 0.70 | 0.84 |
| PC2 (Relative glucosinolate investment) | –0.70 | 0.71 | 0.16 |

**Table B5.** Percent of total trait variation in *Alliaria petiolata* predicted by family, treatment and family-by-treatment interactions after controlling for transplant size. Table demonstrates the effect of including transplant size as a covariate. Only variables where transplant size was a significant covariate are shown.

| **Variables** | **Family** | **Treatment** | **Family × Treatment** |
|---|---|---|---|
| | % ($\chi^2$, p) | % ($\chi^2$, p) | % ($\chi^2$, p) |
| Relative growth rate | 1.89 (3.2, 0.07) | **11.19 (48.9, < 0.001)** | 3.22 (1.3, 0.25) |
| Relative growth rate† | 1.16 (1.3, 026) | **13.74 (60.1, < 0.001)** | 1.68 (0.4, 0.53) |
| TLA | 0.17 (0.9, 0.35) | **27.40 (141.0, < 0.001)** | 2.85 (1.8, 0.19) |
| TLA † | < 0.1 (0, 1) | **28.71 (149.0, < 0.001)** | 2.15 (1.5, 0.22) |
| Fecundity* | < 0.1 (0, 1) | **11.52 (12.0, < 0.001)** | < 0.1 (0, 1) |
| Fecundity*† | 1.08 (0.2, 0.66) | **13.20 (14.56, < 0.001)** | 2.19 (0.1, 0.83) |

Significant estimates ($p < 0.05$) are bolded.

* Measured on log scale

† Transplant size included

**Table B6.** Significance of traits on two performance metrics of *Acer saccharum* (total leaf area in year 1 and shoot mass in year 2), based on likelihood ratio tests. Separate models are performed for treatment effects (i.e. interspecific vs alone) or competitor effects (i.e. interspecific vs treatment).

| **Trait** | **Total leaf area (Year 1)** | **Shoot mass (Year 2)** | **Data in model** |
|---|---|---|---|
| | ($\chi^2$, p) | ($\chi^2$, p) | |
| Treatment | **(17.5, < 0.001)** | **(17.6, < 0.001)** | Interspecific and alone |
| Transplant size | **(150.9, < 0.001)** | **(52.3, < 0.001)** | `` |
| Leaf damage | **(24.1, < 0.001)** | (0.1, 0.8) | `` |
| Fern | **(6.2, < 0.05)** | (0.1, 0.73) | `` |
| Greenhouse bench* | (0, 1) | (0, 1) | `` |
| PC1† | **(11.5, < 0.001)** | (0.4, 0.51) | Interspecific only |
| PC2† | **(5.8, < 0.05)** | (0.7, 0.41) | `` |
| Relative growth rate† | (0, 0.99) | (0, 1) | `` |
| TLA† | (0, 0.89) | **(11.5, < 0.001)** | `` |
| Family† | (20.3, 0.57) | (26.5, 0.23) | `` |
| Transplant size | **(102, < 0.001)** | **(40.9, < 0.001)** | `` |
| Leaf Damage | **(12.5, < 0.001)** | (0, 0.89) | `` |
| Fern | (1.5, 0.22) | (0.3, 0.61) | `` |
| Greenhouse bench* | (0.3, 0.57) | (0, 0.84) | `` |
| Field Column* | NA | (0, 1) | `` |
| Field Row* | NA | **(4.6, < 0.05)** | `` |

Significant estimates ($p < 0.05$) are in bold.

* Random effect

† Competitor trait

**Table B7.** Significance of the effects of *Alliaria petiolata* traits on four different performance metrics ($TLA_{AP}$, survival, fecundity, and lifetime fitness), based on likelihood ratio tests.

| **Competitor/Pathogen trait** | **TLA** | **Survival** | **Fecundity** | **Lifetime fitness** |
|---|---|---|---|---|
| | ($\chi^2$, p) | ($\chi^2$, p) | ($\chi^2$, p) | ($\chi^2$, p) |
| Treatment × PC1 | **(11.3, < 0.01)** | **(7.2, < 0.05)** | (1.1, 0.57) | (4.4, 0.11) |
| Treatment × PC2 | (1.1, 0.57) | (1.8, 0.4) | (2, 0.37) | (0.9, 0.65) |
| Treatment | NA | NA | **(28.5, < 0.001)** | **(17.9, < 0.001)** |
| PC1 | NA | NA | **(4.9, < 0.05)** | (1.6, 0.2) |
| PC2 | (0.4, 0.52) | (2.1, 0.15) | (0.9, 0.35) | (1.2, 0.27) |
| Transplant size | **(11.7, < 0.001)** | (0.4, 0.55) | **(11.8, < 0.001)** | (1, 0.31) |
| Relative growth rate | (0, 0.99) | (0, 1) | (0, 1) | (0, 1) |
| TLA | NA | (2.3, 0.13) | (0.6, 0.45) | (1.6, 0.21) |
| Bolt size | NA | NA | NA | NA |
| Fern | **(5.4, < 0.05)** | **(15.6, < 0.001)** | (0, 0.84) | **(6.7, < 0.01)** |
| Thrip damage | (0.8, 0.37) | (0.3, 0.62) | (0.9, 0.35) | (0.2, 0.66) |
| *Xanthomonas campestris* | **(20.6, < 0.001)** | (0, 0.95) | (0.2, 0.68) | (0.3, 0.57) |
| *Erysiphe cruciferarum* | (1.0, 0.31) | (0.1, 0.72) | (0.8, 0.36) | (0.7, 0.41) |
| Greenhouse bench* | **(38.8, < 0.001)** | (0, 0.89) | (0, 1) | (0, 1) |
| Field column* | NA | (0, 1) | (0, 1) | (0, 1) |
| Field row* | NA | **(7.8, < 0.01)** | (0.3, 0.56) | **(5.0, < 0.05)** |
| Family* | (0.2, 0.69) | **(13.3, < 0.001)** | (0.4, 0.54) | **(14.8, < 0.001)** |

Significant estimates ($p < 0.05$) are in bold.

* Random effect

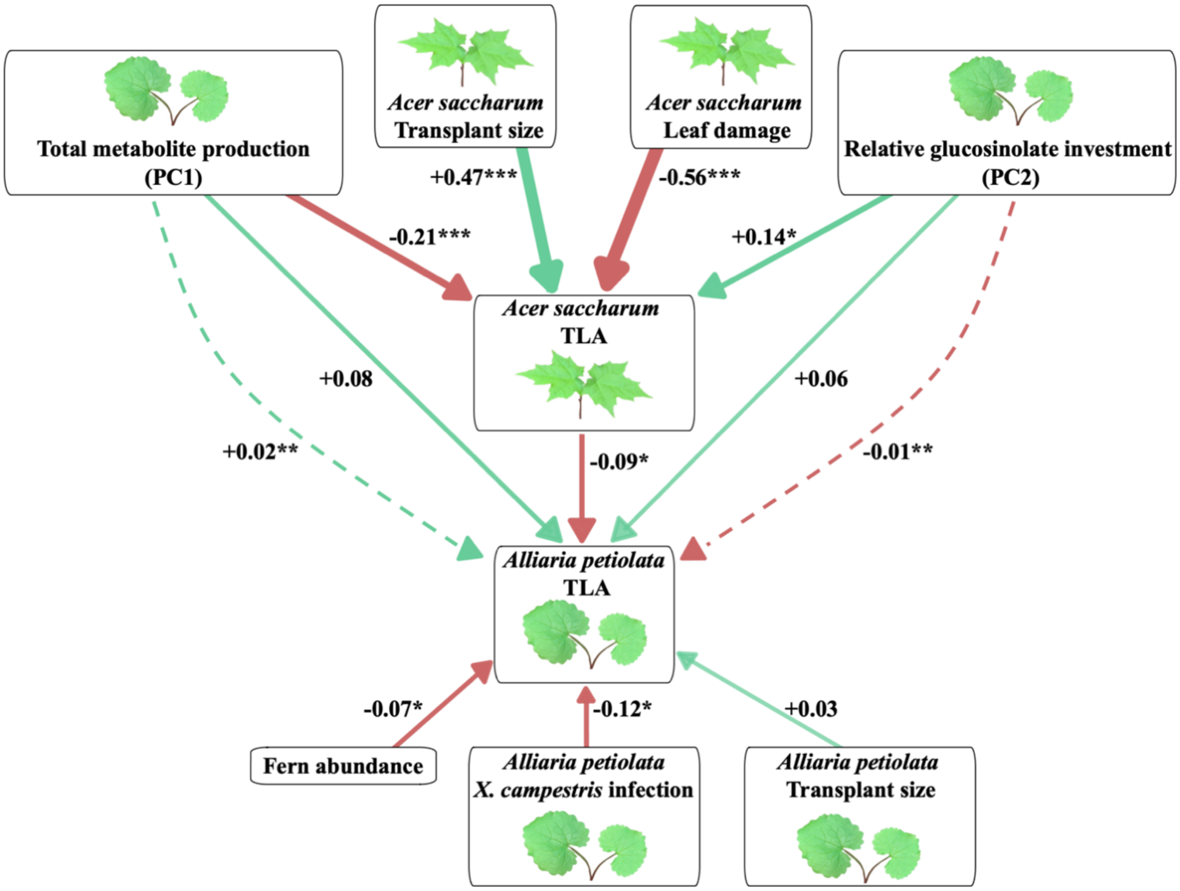


**Figure B1.** Path analysis of *Alliaria petiolata* total leaf area (TLA). TLA was mean-standardized while other variables were z-standardized such that coefficients represent selection gradients. Arrows denote the direction of the effect, with green arrows indicating a positive gradient and red arrows indicate a negative gradient. Arrow thickness denotes the magnitude of the effect and darker arrows indicate greater significance. The text next to each arrow shows the path coefficient and its significance level ($p < 0.05$*, $p < 0.01$**, $p < 0.001$***). Total metabolite production denotes the production of both glucosinolates and Chl *a*, while relative glucosinolate investment indicates the production of glucosinolates relative to Chl *a*. *Acer saccharum* TLA denotes the total leaf area of *Acer saccharum* at the end of year 1. Solid arrows represent direct effects while dashed arrows represent indirect effects.

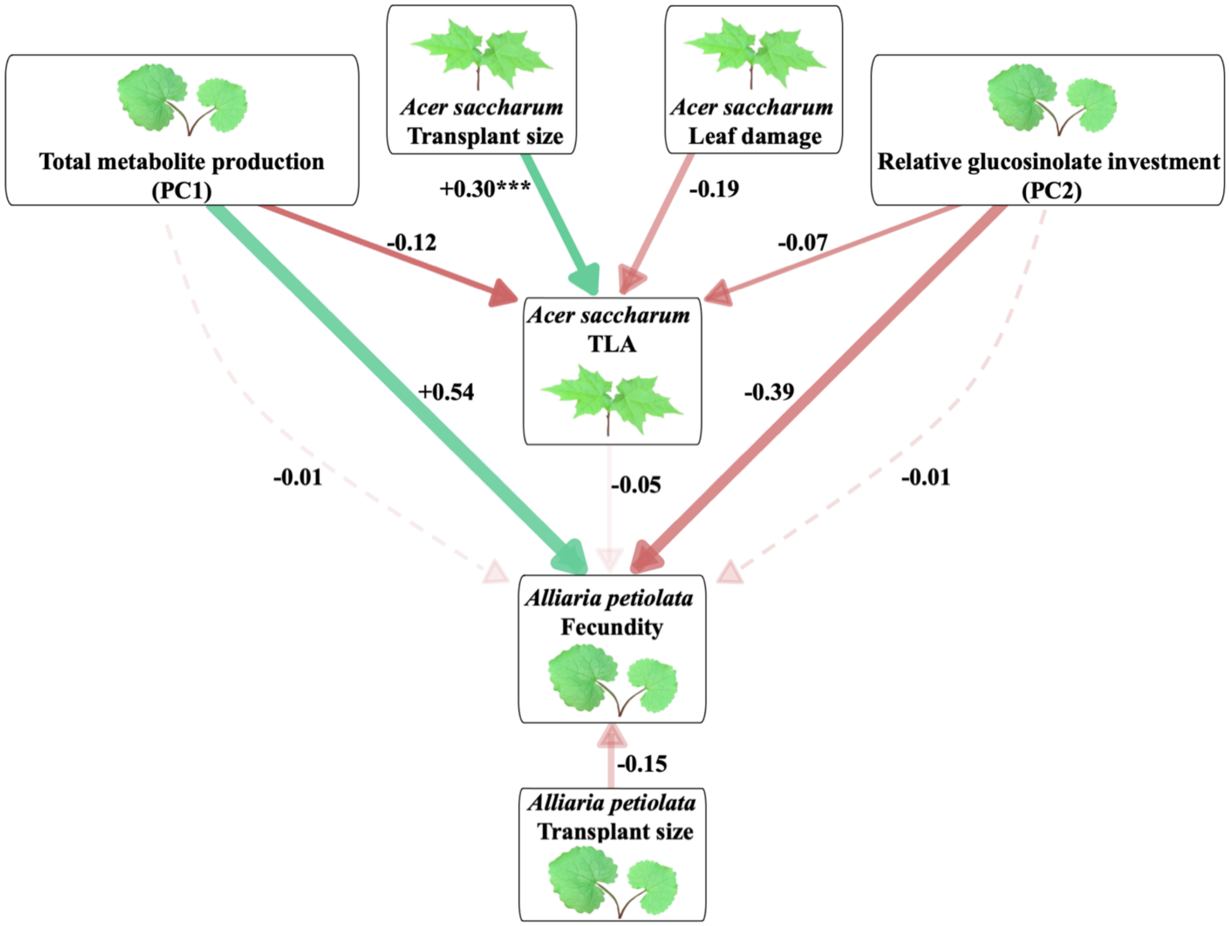


**Figure B2.** Path analysis of *Alliaria petiolata* fecundity. Fecundity was mean-standardized while other variables were z-standardized such that coefficients represent selection gradients. Arrows denote the direction of the effect, with green arrows indicating a positive gradient and red arrows indicate a negative gradient. Arrow thickness denotes the magnitude of the effect and darker arrows indicate greater significance. The text next to each arrow shows the path coefficient and its significance level ($p < 0.05$*, $p < 0.01$**, $p < 0.001$***). Total metabolite production denotes the production of both glucosinolates and Chl *a*, while relative glucosinolate investment indicates the production of glucosinolates relative to Chl *a*. *Acer saccharum* TLA denotes the total leaf area of *Acer saccharum* at the end of year 1. Solid arrows represent direct effects while dashed arrows represent indirect effects.
